# MultiPLIER: a transfer learning framework for transcriptomics reveals systemic features of rare disease

**DOI:** 10.1101/395947

**Authors:** Jaclyn N. Taroni, Peter C. Grayson, Qiwen Hu, Sean Eddy, Matthias Kretzler, Peter A. Merkel, Casey S. Greene

**Affiliations:** Systems Pharmacology and Translational Therapeutics, University of Pennsylvania, Philadelphia, PA, USA; Childhood Cancer Data Lab, Alex’s Lemonade Stand Foundation, Philadelphia, PA, USA; National Institute of Arthritis and Musculoskeletal and Skin Diseases, National Institutes of Health, Bethesda, MD, USA; Division of Nephrology, Department of Internal Medicine, Michigan Medicine, Ann Arbor, MI, USA; Department of Computational Medicine and Bioinformatics, Michigan Medicine, Ann Arbor, MI, USA; Division of Rheumatology and the Department of Biostatistics, Epidemiology, and Informatics, University of Pennsylvania, Philadelphia, PA, USA; Institute of Translational Medicine and Therapeutics, University of Pennsylvania, Philadelphia, PA, USA; Institute of Biomedical Informatics, University of Pennsylvania, Philadelphia, PA, USA

## Abstract

Unsupervised machine learning methods provide a promising means to analyze and interpret large datasets. However, most gene expression datasets generated by individual researchers remain too small to fully benefit from these methods. In the case of rare diseases, there may be too few cases available, even when multiple studies are combined. We trained a Pathway Level Information ExtractoR (PLIER) model using on a large public data compendium comprised of multiple experiments, tissues, and biological conditions. We then transferred the model to small rare disease datasets in an approach we term MultiPLIER. Models constructed from large, diverse public data i) included features that aligned well to important biological factors; ii) were more comprehensive than those constructed from individual datasets or conditions; iii) transferred to rare disease datasets where the models describe biological processes related to disease severity more effectively than models trained on specifically those datasets.

## INTRODUCTION

The rapid expansion of the amount of publicly available gene expression data presents opportunities for discovery-driven research into rare diseases with poorly understood etiologies. Methods that extract a low-dimensional representation of the genome-scale gene expression data are useful for biological discovery, particularly in a multi-dataset setting. Such methods are biologically motivated, as genes that vary together in their expression may function together (Myers et al., 2006). There are also technical considerations, as measuring combinations of genes reduces the multiple hypothesis testing burden, can aid feature engineering, and is more likely to yield robust results than analyses of individual gene measurements (Cleary et al., 2017). These methods have advantages over two-group gene set-based comparisons because they provide more context for genes, are better fit to the underlying data, and remove the difficulty of identifying the most useful comparisons *a priori* (Stein-O’Brien et al., 2017a). Unsupervised machine learning (ML) methods including matrix factorization- and autoencoder-based approaches have successfully extracted biologically meaningful low-dimensional representations of gene expression data that can distinguish disease types, predict drug response, and identify new pathway regulators (Dincer et al., 2018; Stein-O’Brien et al., 2017b; Tan et al., 2017; Way and Greene, 2017).

Studying rare diseases using this data-intensive methodology is challenging. Even if multiple datasets exist for a disease of interest, the experimental designs may not be comparable or controls may not be selected by the same means. Combining a longitudinal clinical trial that lacks healthy controls and an untreated case-control study may not be appropriate. One strategy to integrate multiple, distinct datasets that cannot be directly combined is to identify modules or factors within each dataset and to seek to identify common changes within the set of identified patterns. A challenge with this approach is that the factors, though they may capture similar processes, are not identical. Especially in the case where a one-to-one mapping for factors across datasets is lacking, this strategy can require human examination. This can be particularly time consuming because the number of comparisons grows with the square of the number of datasets.

Training a single, comprehensive model and using the same model for the analysis of datasets of interests allows the same factors to be directly compared. Moreover, signals may vary together in a particular small disease dataset but in fact be representative of distinct processes (e.g., glycolysis and an increase in infiltration of a specific immune cell type or change in tissue composition). Models trained on these data alone may not distinguish correlated processes. However, with a large, uniformly processed data source comprised of many conditions, including those where these processes are effectively “decoupled” (e.g., cell line experiments where glycolysis is perturbed but immune infiltration is irrelevant), it may be feasible to derive features from the large, heterogeneous compendium and transfer them to rare disease datasets. This is an example of *unsupervised transfer learning* and “feature-representation-transfer” (Pan and Yang, 2010). Intuitively, we might expect the features constructed from training on the data from heterogeneous conditions to be more comprehensive and more broadly applicable to various biological conditions and experimental designs.

We wanted to determine if knowledge learned in a source domain—here, a large collection of human gene expression data—could be transferred to a target domain to improve the target task performance—here, the detection of patterns that characterize disease in unseen disease states. In structured data, such as the LINCS L1000 expression data (Subramanian et al., 2017), using information from *similar conditions* (e.g., drugs and cell types) to predict expression profiles has been shown to be useful for downstream machine learning tasks (Hodos et al., 2017). We sought to extract a biologically meaningful low-dimensional representation of the recount2 compendium—a uniformly processed human RNA-seq dataset comprised of tens of thousands of samples and thousands of experiments (Collado-Torres et al., 2017)—and project new datasets into this low-dimensional representation for downstream analysis. In contrast to the L1000 example, recount2 is less specific to the task at hand and represents a generic collection of human samples assayed by the scientific community at large. For knowledge transfer to be successful, the domains must be sufficiently related. We posited that transcriptional regulation, the resulting modular nature of gene expression data, and molecular mechanisms shared among diseases would support unsupervised transfer learning.

Just as the similarity between domains is important, selecting an appropriate method is crucial. We used the Pathway-Level Information ExtractoR (PLIER) matrix factorization framework (Mao et al., 2017). PLIER leverages prior biological knowledge (supplied as gene sets to the model), but can also identify features not associated with these sets. It learns patterns of correlated genes, termed latent variables (LVs). In this case the latent variables are the low-dimensional representations. PLIER imposes certain constraints such that some of the learned latent variables will align with the input gene sets. Because not *all* latent variables are forced to relate to gene sets, some will capture major sources of variation due to technical factors. PLIER was shown to be robust to expression data processing choices, particularly for gene set-associated latent variables (Mao et al., 2017), though it was not used in the context of a multi-dataset compendium. PLIER is well-suited for our task because capturing *shared biology* is key to transfer learning and because there will necessarily be technical noise in the recount2 compendium. We call this approach and the downstream analysis MultiPLIER because we train the model on multiple tissues and diseases. We demonstrate that MultiPLIER is appropriate for biological discovery in multiple contexts, including cross-platform (microarray) applications.

To evaluate MultiPLIER, we designed experiments to assess the recovery of biologically meaningful latent variables, evaluate how training set characteristics such as sample size influence what is captured by PLIER models, and measure the concordance between MultiPLIER and disease- and dataset-specific models. We center our downstream analysis on antineutrophil cytoplasmic autoantibody (ANCA)-associated vasculitis (AAV), a rare systemic autoimmune disorder, because it affects multiple tissues, too few gene expression assays exist to support data-intensive unsupervised machine learning, and the underlying molecular mechanisms are not fully understood. We demonstrate that PLIER learns cell type-specific signatures and reduces technical batch effects when trained on a multi-dataset microarray compendium of systemic lupus erythematosus (SLE) whole blood (WB). We observe that MultiPLIER, trained on recount2, retains the biological fidelity of a compendium-specific model in the SLE whole blood evaluations. We find that increasing sample size and breadth of biological conditions included in training sets yielded models that captured more biological pathways and distinguished similar molecular processes, supporting the use of our approach. MultiPLIER is concordant with dataset-specific models in AAV and also identifies altered molecular processes in severe or active disease in multiple tissues. Examining two medulloblastoma cohorts with the same MultiPLIER model reveals consistent pathway differences across subtypes, demonstrating its applicability to different classes of rare diseases. In summary, MultiPLIER’s features have biological relevance and can be directly compared across datasets making it particularly valuable for the integrative analysis of complex human disease.

## RESULTS

### The MultiPLIER framework for unsupervised transfer learning

PLIER can be applied to or trained on individual datasets or large compendia (Mao et al., 2017). In Figure 1, we illustrate two different approaches to analyzing a multisystem disease with PLIER. PLIER automatically extracts latent variables, shown as the matrix *B*, and their loadings (*Z*) (Figure 1A). We refer to the entire collection of latent variables (*B*) as the latent space. In this example, three datasets from the same complex multisystem disease and distinct tissues are each used as a training set for a PLIER model. We refer to these PLIER models as *<u>disease-</u> <u>specific</u> <u>or</u> <u>single</u> <u>dataset</u> <u>models</u>*. This training approach results in three dataset-specific latent spaces (Figure 1A). We provide the following information on the PLIER method to provide background for our choice of model and illustrate the potential benefits of training on a large, diverse compendium.

**Figure 1.**
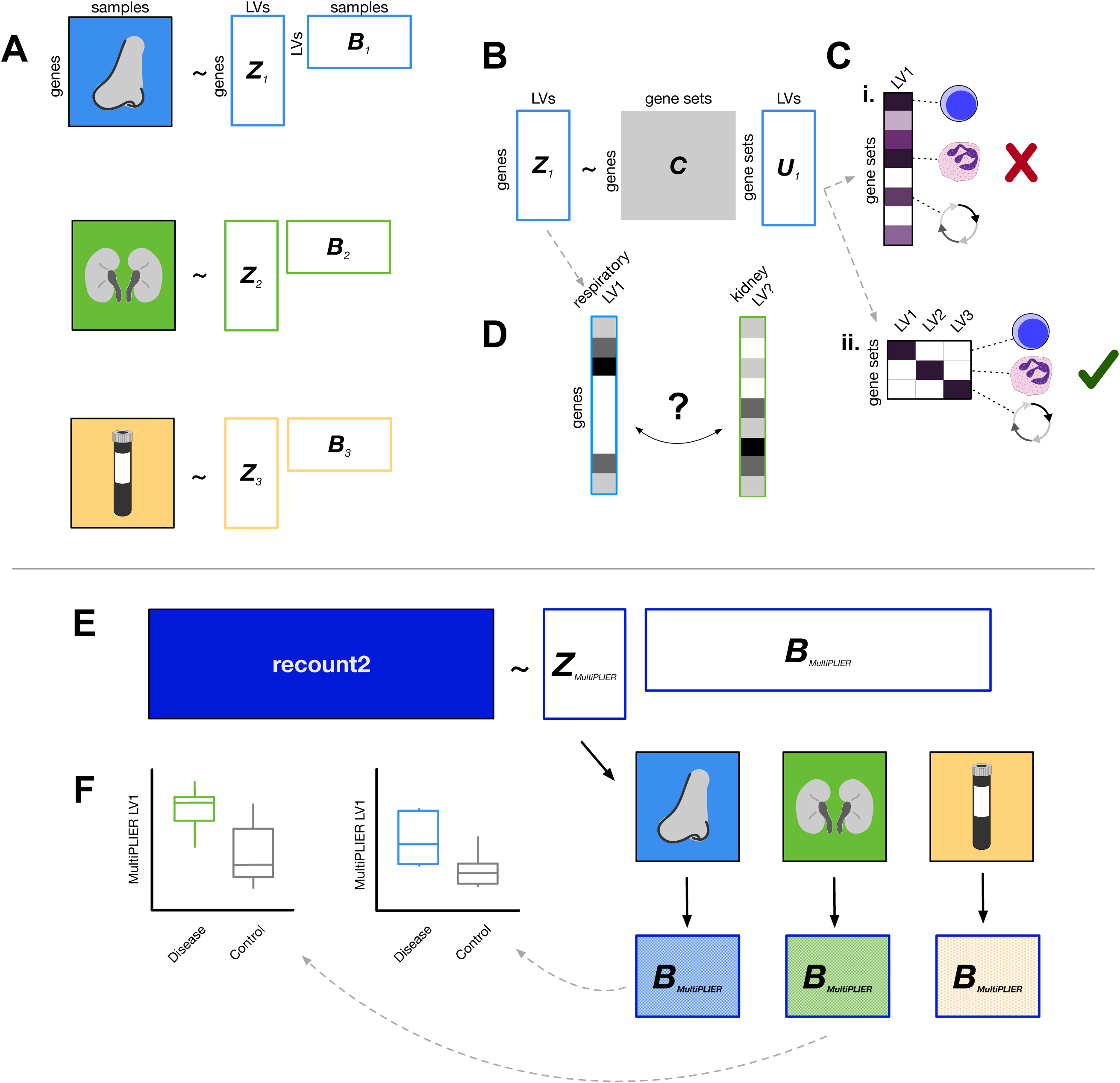
Overview of dataset-specific PLIER and MultiPLIER. Boxes with solid colored fills represent inputs to the model. White boxes with colored outlines represent model output. **(A)** PLIER (Mao et al., 2017) automatically extracts latent variables (LVs), shown as the matrix *B*, and their loadings (*Z*). We can train PLIER model for each of three datasets from different tissues, which results in three dataset-specific latent spaces. **(B)** PLIER takes as input a prior information/knowledge matrix *C* and applies a constraint such that some of the loadings (*Z*) and therefore some of the latent variables capture biological signal in the form of curated pathways or cell type-specific gene sets. **(C)** Ideally, a latent variable will map to a single gene set or a group of highly related gene sets to allow for easy interpretation of the model. PLIER applies a penalty on *U* to facilitate this. Purple fill in a cell indicates a non-zero value and a darker purple indicates a higher value. We show an undesirable *U* matrix in the top toy example (Ci) and a favorable *U* matrix in the bottom toy example (Cii). **(D)** If models have been trained on individual datasets, we may be required to find “matching” latent variables in different dataset- or tissue-specific models using the loadings (*Z*) from each model. Using a metric like the Pearson correlation between loadings, we may or may not be able to find a well-correlated match between datasets. **(E)** The MultiPLIER approach: train a PLIER on a large collection of uniformly processed data from many different biological contexts and conditions (recount2; Collado-Torres et al., 2017)—a MultiPLIER model—and then project the individual datasets into the MultiPLIER latent space. The hatched fill indicates the sample dataset of origin. **(F)** Latent variables from the MultiPLIER model can be tested for differential expression between disease and controls in multiple tissues.

PLIER constrains the loadings *Z* to align with curated pathway or gene expression-derived cell type-specific gene sets, specified in the prior knowledge matrix *C* which is input to the model (Figure 1B). This ensures that some but not all of the resulting latent variables capture *known biology* and is an important and attractive feature of this particular approach. The prior information coefficient matrix *U* captures the relationships between latent variables and the gene sets. In this example, the PLIER model is trained using the same input gene sets.

PLIER applies a penalty to *U* such that an individual latent variable should represent only a few gene sets (Figure 1C). The penalty makes latent variables more interpretable. Ideally, an individual latent variable will be unambiguously associated with one pathway or cell type gene set or a collection of closely related gene sets. In Figure 1C, we show hypothetical latent variables (columns) from the *U* matrix. In the top example, LV1 is strongly associated with lymphocyte, granulocyte, and cell cycle gene sets (Figure 1Ci). This association with many gene sets is undesirable because it hampers our ability to interpret the latent variable. Training on small datasets is more likely to result in latent variables that are associated with multiple pathways or cell types because there are fewer samples to disambiguate confounded factors. For instance, if neutrophil and T lymphocyte composition of all disease samples is increased relative to controls in a small dataset, these two signals may be difficult to distinguish. In the bottom example, latent variables are each associated with a single pathway or cell type. In the bottom toy example, latent variables are each associated with a single pathway or cell type—the ideal case (Figure 1Cii). PLIER additionally calculates AUC and p-value metrics for the association between a gene set and a latent variable.

In the case of multisystem disorders, identifying molecular processes that are perturbed across tissues is often desirable and may shed light on the pathobiology of that disease. In the dataset-specific PLIER case, we must map between individual dataset models—finding latent variables that are similar to one another (Figure 1D). This can be challenging, as not all genes are represented in different datasets, multiple biological signals may be captured by a single latent variable if they tend to vary together (i.e., the neutrophil-T lymphocyte example above). Thus, a latent variable may or may not have a match across datasets, limiting the multi-tissue analyses that can be performed downstream.

An alternative to the disease-specific or single-dataset PLIER approaches is to train on a large collection of uniformly processed data from numerous biological contexts and conditions and then to use the loadings (*Z*) from this model to project the individual datasets into the *same latent space* (Figure 1E). This is the approach we term *<u>MultiPLIER</u>*. In this work, we use the subset of the recount2 compendium (Collado-Torres et al., 2017) that includes metadata as our training set. This RNA-seq training set is comprised of approximately 37,000 samples, less than 100 of which are predicted to be from autoimmune disease and the majority of which are labeled as cell lines or tissue samples by the MetaSRA project for normalization of RNA-seq sample metadata (Bernstein et al., 2017). We refer to this model as either MultiPLIER model or the full recount2 model to signify its training set and to latent variable 1 from this model as MultiPLIER LV1. The large sample size and breadth of molecular processes represented increases the number of gene sets that the model captures and the *specificity* of the model—the learned latent variables are more interpretable due to a “separation” of signals.

Once the individual datasets are in the same latent space, the latent variables can serve as input into numerous downstream applications, including supervised tasks such as predicting response to treatment or testing for differential expression. In the Figure 1F toy example, MultiPLIER-learned latent variables showed the same directionality (increased expression) in multiple tissues. This suggests that these molecular processes are systematically altered and warrants further investigation. For more information on implementation, see STAR Methods and the PLIER preprint (Mao et al., 2017).

### A PLIER model trained on a systemic lupus erythematosus whole blood compendium learned SLE pathology-relevant latent variables and differentiated biological signal from technical noise

We sought to determine if PLIER was appropriate for use with compendia assembled from multiple publicly available datasets. We used SLE whole blood as a case study because of the molecular heterogeneity of the disease that is apparent at the gene expression (Banchereau et al., 2016) and because whole blood is a mixture of immune cells; this is a condition where we expected both the cell type composition and phenotype to be altered. We assembled an SLE whole blood compendium from 7 publicly available datasets that used different microarray platforms (Table 1). We selected these datasets because they included specific perturbations (e.g., treatments with a targeted biologic) or sample metadata (e.g., cell type counts) that supported evaluations of the model. In addition, some datasets had only submitter-processed data available, which can occur in repositories like Gene Expression Omnibus and ArrayExpress (Barrett et al., 2012; Edgar, 2002; Kolesnikov et al., 2014). We assembled the compendium by uniformly processing available raw data and scaling individual datasets (see STAR Methods for details). Despite this processing, dataset-specific factors were still evident in the first five principal components (PCs) (Figure S1).

**Table 1.** Datasets included in the systemic lupus erythematosus whole blood dataset. Raw data availability refers to whether or not raw data was associated with the accession on ArrayExpress. We note any information that is relevant to the evaluations we performed.

| Accession | Number of samples | Platform | Raw data availability | Publication |
| --- | --- | --- | --- | --- |
| E-GEOD-65391* | 996 | Illumina HumanHT-12 v4 | No | Banchereau et al., 2016 |
| E-GEOD-11907 | 4 | Affymetrix HGU133A and B | Yes | Chaussabel et al., 2008 |
| E-GEOD-39088** | 142 | Affymetrix HGU133plus2 | Yes | Lauwerys et al., 2013 |
| E-GEOD-72747 | 30 | Affymetrix HGU133plus2 | Yes | Ducreux et al., 2016 |
| E-GEOD-49454 | 177 | Illumina HumanHT-12 v4 | No | Chiche et al., 2014 |
| E-GEOD-61635 | 129 | Affymetrix HGU133plus2 | Yes | N/A |
| E-GEOD-78193*** | 125 | Agilent 4x44K | No | Welcher et al., 2015 |
| *Notes: \*Longitudinal pediatric cohort; Accession includes neutrophil counts \*\*IFN-alpha-kinoid trial \*\*\*AMG 811 trial* |  |  |  |  |

We trained a PLIER model on this SLE WB compendium (termed the SLE WB PLIER model) and analyzed the learned latent variables. Many cell types found in whole blood were represented (CD4 T cells in LV86, Monocytes in LV72 and LV109), as well as latent variables associated with type I IFN signaling (LV6 and LV69) and highly relevant to autoimmunity like KEGG Graft vs. Host Disease and KEGG Antigen processing and presentation (LV59) (Figure 2A). Since PLIER does not require all learned latent variables to be associated with gene sets, it can also learn major sources of variation associated with technical noise. As noted, the SLE WB compendium showed evidence of technical noise following processing (Figure S1).

**Figure 2.**
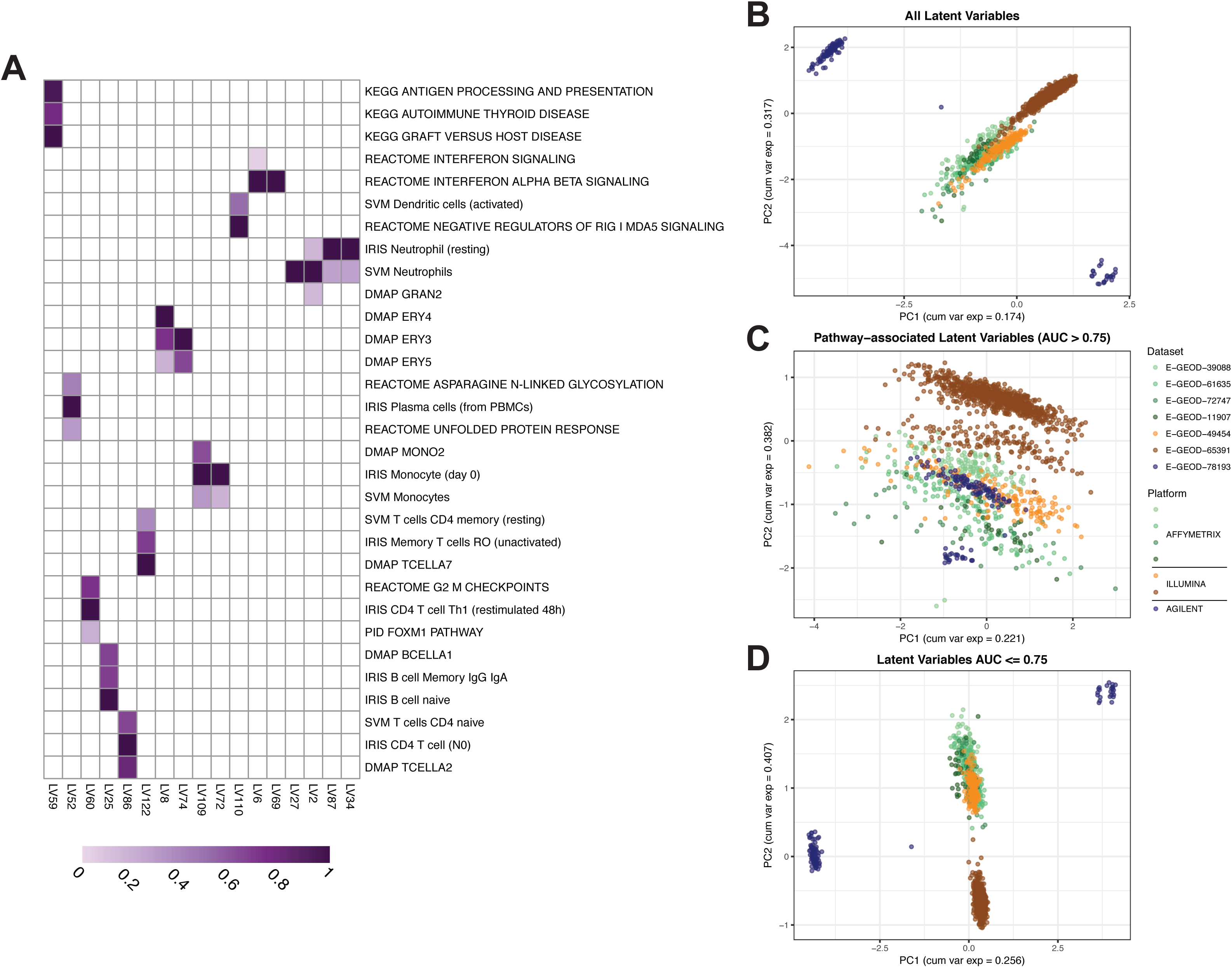
A PLIER model trained on a systemic lupus erythematosus (SLE) whole blood (WB) compendium learns SLE pathology-relevant latent variables and divides biological signal and technical noise. **(A)** Selected latent variables (LVs) from the SLE WB PLIER *U* matrix. Purple fill in a cell indicates a non-zero value and a darker purple indicates a higher value. Only pathways with AUC > 0.75 in displayed latent variables are shown. Panels B-D display the first two PCs from different subsets of the latent space or *B* matrix. Points are samples. Samples are colored by dataset of origin and datasets from the same platform manufacturer are similar colors. **(B)** PC1 and PC2 from PCA on the entire B matrix illustrates a platform- or dataset-specific effect. **(C)** PC1 and PC2 from only pathway- or geneset-associated latent variables (LVs with AUC > 0.75 for at least one geneset) show a reduction in the technical variance evident in panel B. **(D)** PC1 and PC2 from only latent variables that do not have an association with an input geneset (all AUC <= 0.75) show a similar pattern to that of all latent variables. The dataset-specific effect in panels B and D is also observed in PCA on the gene-level gene expression data and at different AUC thresholds (Fig. S1).

We hypothesized that latent variables associated with biological signals would exhibit less technical noise. We performed principal component analysis (PCA) on different subsets of the latent space or *B* matrix for visualization purposes. With PCA on all latent variables (Figure 2B), we observe separation of the Agilent dataset (E-GEOD-78193) in PC1 and PC2. Once the B matrix was subset to only latent variables associated with pathways or cell types (AUC > 0.75 for at least one gene set), less separation between datasets was observed (Figure 2C). Note that E-GEOD-65391 is a pediatric cohort, which may account for some of the separation in PC2 in Figure 2C. In contrast, when examining only latent variables that are not explicitly tied to gene sets (Figure 2D), the dataset-specific effect was more exaggerated than in all latent variables (Figure 2B). This was observed for a variety of AUC cut points for pathway-association (Figure S1B-G).

As illustrated in Figure 2A, PLIER learned cell type-specific signals from heterogeneous expression data. PLIER was also shown to outperform other methods (e.g., CIBERSORT, Newman et al., 2015; non-negative matrix factorization) in estimating cell type proportions from a small dataset of whole blood RNA-seq samples as measured by correlation to counts from mass cytometry data (Mao et al., 2017). We sought to evaluate whether or not these signals captured in the SLE WB model were well-correlated with cell type counts, despite the fact that the model was not explicitly trained to predict cell type composition. We used the Banchereau, et al. dataset (E-GEOD-65391) from the SLE WB compendium which includes neutrophil counts in the sample metadata for this evaluation. We selected LV87 from the SLE WB PLIER model (SLE WB LV87) from multiple latent variables that were significantly associated (FDR < 0.05) with neutrophil gene sets because it lacked an association with other myeloid cell types (e.g., monocyte- or macrophage-related gene sets) and could then be considered to be specific to neutrophil gene sets. We observed that SLE WB LV87 showed agreement (*R*^2^ = 0.29) with the E-GEOD-65391 neutrophil counts (Figure 3A).

**Figure 3.**
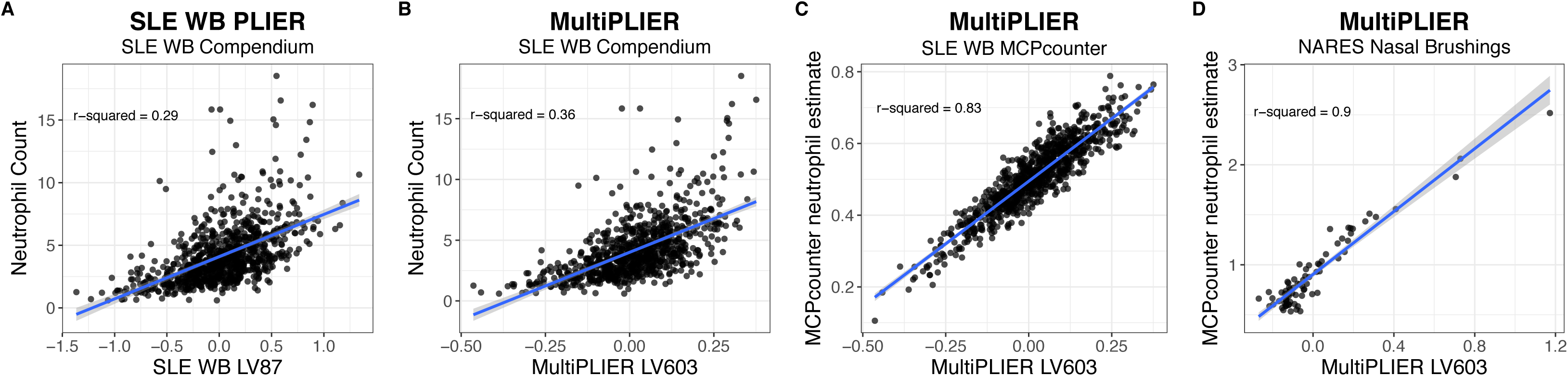
MultiPLIER learns a neutrophil-associated latent variable (LV) that is well-correlated with neutrophil counts or estimates in multiple tissues and disease contexts. **(A)** SLE WB LV87, LV87 from the PLIER model trained on the entire SLE WB compendium, can predict neutrophil count in Banchereau, et al. dataset (Banchereau et al., 2016) despite being entirely unsupervised with respect to this task. **(B)** The MultiPLIER neutrophil-associated latent variable (LV603) performs slightly better than the SLE WB-specific model. Here, the SLE WB compendium is projected into the MultiPLIER latent space. Recall that the MultiPLIER model has not been exposed to the *specific* technical variance found in the SLE WB compendium. **(C)** MultiPLIER LV603 values are highly correlated from a state-of-the-art method for estimating immune infiltrate, MCPcounter (Becht et al., 2016). This suggests that the modest correlation with neutrophil count in B is the result of estimating neutrophil, a terminally differentiated cell type, counts from transcriptome data, rather than a limitation of the MultiPLIER approach. **(D)** MultiPLIER performance is not limited to whole blood, as it is highly correlated with the MCPcounter neutrophil estimate in the NARES dataset (Grayson et al., 2015). NARES is a nasal brushing microarray dataset that includes patients with ANCA-associated vasculitis, patients with sarcoidosis and healthy controls among other groups and was projected into the MultiPLIER latent space.

### Training on a larger and more heterogeneous compendium yielded more comprehensive models

In SLE, we established the feasibility of applying PLIER to a multi-dataset compendium comprised of a mixture of cell types. However, not all pathways that are potentially relevant to SLE were captured by the SLE WB PLIER model (e.g., type II interferon signaling; Chiche, et al. 2014.). We evaluated the extent to which a dataset’s sample size or breadth contributed to desirable characteristics of PLIER models—namely, the representation of more pathways. We performed two experiments to assess the effect of sample size and training set composition, respectively: 1) randomly selecting *n* samples for a range of sample sizes and 2) subsetting recount2 using terms from biomedical ontologies as labeled by MetaSRA (Bernstein et al., 2017). Specifically, we used training sets comprised of blood samples, cancer samples, tissue samples, cell line samples, and samples that were not labeled as blood samples, termed “other tissues” (see STAR Methods for more information on the ontologies and terms). We performed 5 repeats for each evaluation, where a different set of *n* samples was included in training (sample size) or the same training set was used but the model was initialized with different random seeds (biological context). We evaluated models by the following metrics: the number of latent variables (*k*), the proportion of input gene sets with at least one latent variable significantly associated (FDR < 0.05; termed “pathway coverage”), and the proportion of latent variables that were significantly associated with at least one pathway (FDR < 0.05). The results are shown in Figure 4A-C, where biological contexts are ordered such that they increase in sample size (indicated in Figure 4D) with the full recount2 MultiPLIER model (n = 37032) for comparison.

**Figure 4.**
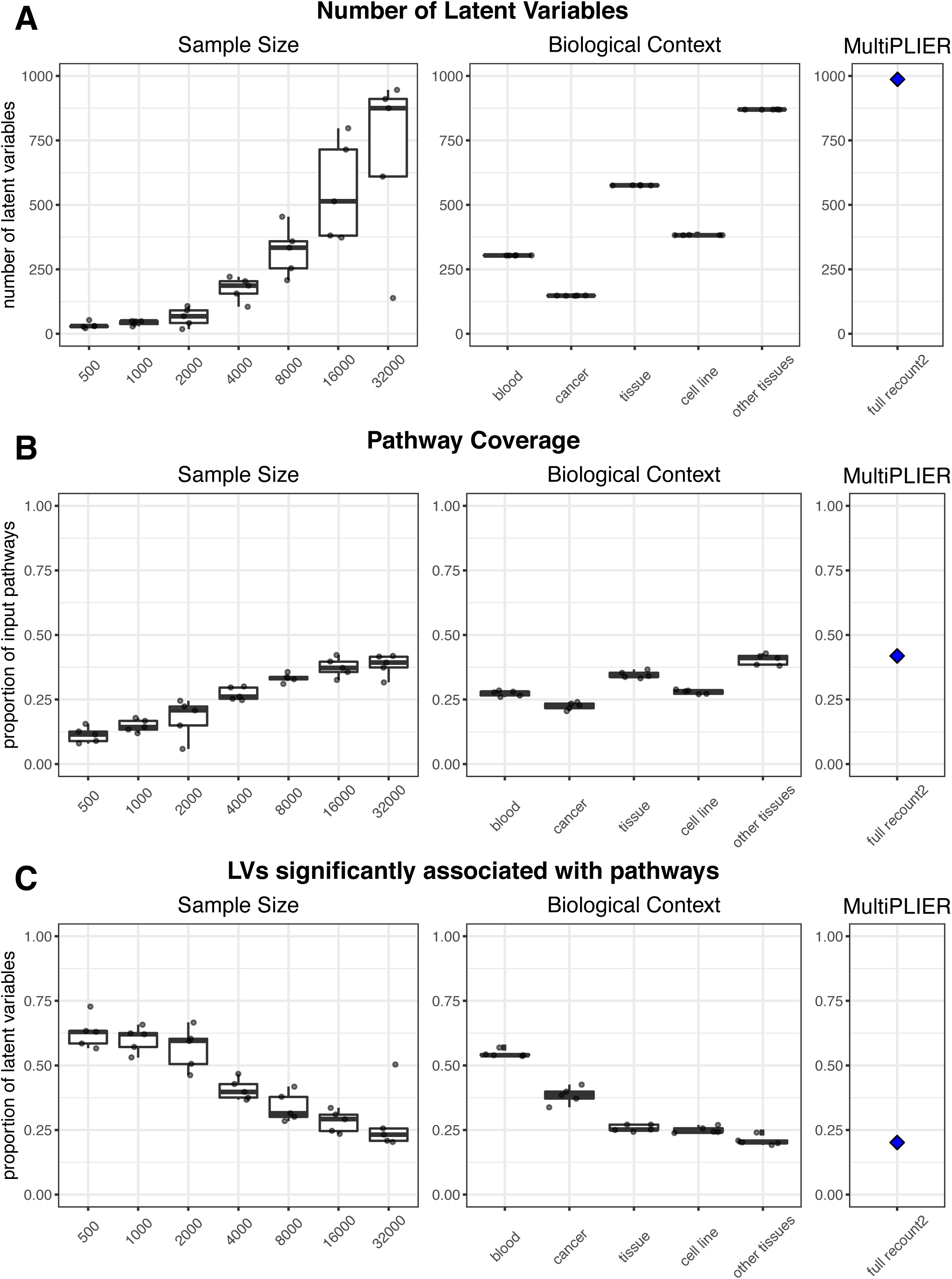
Subsampling of the recount2 compendium demonstrates the contribution of both sample size and breadth of biological conditions to PLIER model characteristics. PLIER models were trained on samples randomly selected from the recount2 compendium (sample size evaluations) or on a subset of the recount2 compendium mapped to the same ontology term in MetaSRA (Bernstein et al., 2017) (biological context evaluations; see STAR Methods for the specific terms used). The training set for each repeat in the biological context evaluations is comprised of the *same samples*, but initialized with different random seeds. The boxplot and points in black in A-C represent 5 repeats performed for each sample size or biological context. The blue diamonds and panels labeled MultiPLIER are the values from the full recount2 PLIER model (∼37,000 samples). The sample size for each biological context training set is below the biological context heatmap in panel D; the biological contexts are ordered by increasing sample size in all panels. **(A)** The number of latent variables (*k*) in a model is generally dependent on sample size. However, the biological contexts where samples are expected to be comprised of a mix of cell types (e.g., blood and tissue) have a high number of latent variable (LV) than we would expect based on the sample size experiments. **(B)** The proportion of pathways supplied as input to the model that are significantly associated (FDR < 0.05) with at least one latent variable, termed pathway coverage, mirrors the number of latent variables in a model. **(C)** The proportion of latent variables that are significantly associated (FDR < 0.05) with at least one pathway or gene set generally decreases with sample size. The exceptions are models trained on blood, which is likely the most homogeneous of the training sets, and many gene sets supplied to the models during training are immune cell related which this training set is well-suited to capture. This suggests that increasing the sample size or breadth of the training set introduces more signal that is not biologically relevant, at least with respect to the pathways that have been supplied to the model.

In general, the larger a training set, the higher *k* (Figure 4A, Sample Size); *k* is set by first determining the number of significant PCs (or major variance components) identified in the dataset. Thus, there tended to be more major variance components in larger datasets. The exceptions to this were the models trained on the two biological contexts where we expected many samples to be of a mixture of various cell types, blood (n = 3862) and tissue (n = 12396), where *k* was higher than expected based on the sample size alone (Figure 4A).

Models with more latent variables had higher pathway coverage in both the sample size and biological context experiments (Figure 4B). In addition, the MultiPLIER model achieved pathway coverage = 0.767 for the MSigDB oncogenic pathways (Subramanian et al., 2005), which were not included during model training. The *identity* of the pathways captured by the model was also influenced by the biological conditions included in the training set. For instance, models trained on blood were uniquely able to capture more specialized or mature immune cell gene sets. Models trained on the cancer training set (n = 8807; note that the cancer training set includes many cell line samples because of the annotation source) were unable to capture natural killer (NK) cell signatures whereas that the random subsampling sets of a similar size (n = 8000) captured this cell type. The proportion of latent variables significantly associated (FDR < 0.05) with at least one pathway generally decreased with sample size (Figure 4C). Models trained on blood were an exception, as this proportion was higher than the random subsampling of a similar sample size (n = 3862 vs. n = 4000 random samples).

### MultiPLIER learned cell type-associated latent variables that are well-correlated with counts or estimates in multiple tissues and disease contexts

The MultiPLIER model captured a large number of biological signals, including oncogenic pathways that were “novel” to the model. We next sought to determine the extent to which models trained on large and heterogeneous datasets would be comparable to individual dataset-specific models or useful for the downstream analysis of small datasets. We projected the SLE WB compendium into the MultiPLIER latent space to compare the MultiPLIER and SLE WB PLIER models. We evaluated neutrophil counts using MultiPLIER LV603, which was significantly associated (FDR < 0.05) with several neutrophil gene sets and not significantly associated with gene sets from other myeloid lineage cell types (Figure S2A). We observed stronger agreement between MultiPLIER LV603 expression and neutrophil counts (*R*^2^ = 0.36) than between SLE WB LV87 and the counts (*R*^2^ = 0.29; Figure 3A-B). The correlation between MultiPLIER LV603 and the neutrophil counts was relatively modest, though this is the highest correlation of any latent variable from the MultiPLIER model. Comparing the MultiPLIER model’s LV603 with neutrophil estimates from MCPcounter (Becht et al., 2016), a method specifically designed for estimating immune cell infiltration, reveals high correlation (*R*^2^ = 0.83; Figure 3C). This suggests that correlation with measured counts is limited because neutrophils are mature terminally differentiated cells.

Agreement with the Banchereau, et al. data was not limited to the neutrophil signatures (Banchereau et al., 2016). In the original publication, the authors stratified patients into three groups based on their SLEDAI (SLE Disease Activity Index) scores (Rovenský and Payer, 2009) and demonstrated that those patients with higher disease activity had increase plasmablast counts (as measured by FACS in a subset of patients). Both the SLE WB PLIER and MultiPLIER plasma cell associated latent variables showed the same pattern as was reported in the original publication (Figure S2B-C) (Banchereau et al., 2016). MultiPLIER, trained exclusively on RNA-seq data, recovered these features even though it was not trained on this dataset and had not encountered the particular technical noise in the SLE WB microarray compendium.

To assess whether or not this performance was generalizable to other datasets or disease conditions (including solid tissues), we performed additional evaluations in a nasal brushing microarray dataset (NARES dataset) that included patients with granulomatosis with polyangiitis (GPA), sarcoidosis, and healthy controls (Grayson et al., 2015) and a microarray dataset of isolated immune cell types from different autoimmune conditions (E-MTAB-2452; McKinney et al., 2015). Neutrophil counts were unavailable for the NARES solid tissue dataset. Instead, we again use MCPcounter neutrophil estimates (Becht et al., 2016), which was demonstrated to work in solid tissues including non-cancerous tissues, as our dependent variable. MultiPLIER LV603 was highly correlated (*R*^2^ = 0.9) with the MCPcounter neutrophil estimate in the NARES dataset (Figure 3D). The isolated leukocyte dataset included samples that were selected for CD4 (T cells), CD14, (monocytes) and CD16 (neutrophils). MultiPLIER latent variables associated with particular cell type gene sets were generally most highly expressed in the samples comprised of that isolated cell type (Figure S3).

### A large and diverse training set allowed PLIER models to distinguish similar pathways

We assessed the effect of training set sample size and composition on a model’s ability to distinguish related pathways (e.g., learn disentangled features), termed “pathway separation,” with the models trained on different biological contexts and across different samples sizes. We considered two similar sets of pathways or gene sets to be successfully separated by a model if at least one pathway or gene set from each set of similar pathways was captured by the model (FDR < 0.05) and each set was uniquely represented. An example using neutrophil- and monocyte/macrophage-related gene sets: we considered a model that had at least one neutrophil-associated latent variable that was not associated with any monocyte/macrophage gene sets and at least one monocyte/macrophage-associated latent variable that was not associated with any of the neutrophil gene sets to have separated these pathways. This experiment is labeled “myeloid” in Figure 5 and was selected because of the shared lineage of these cell types. We also evaluated models’ abilities to separate type I and type II interferon signaling (“IFN”) and the G1 and G2 phases of the cell cycle (“proliferation”). (See STAR Methods and Table 3 for detailed information about the gene sets.) In the sample size and biological context evaluations, we trained 5 models. The color of the heatmap cells in Figure 5 indicates the number of models where we found pathway separation for each pair of gene sets (row) and condition (column).

**Figure 5.**
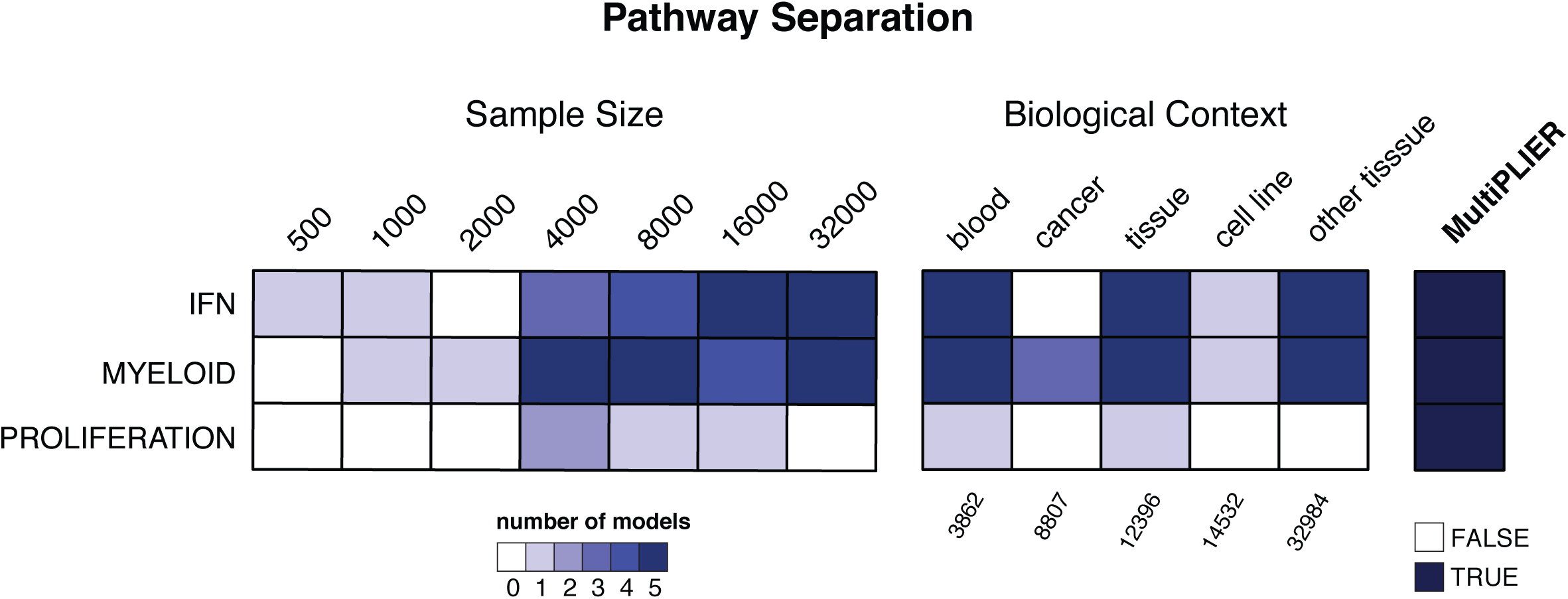
MultiPLIER distinguishes related pathways; both the sample size of and biological conditions within a training set affect a model’s ability to do so. Pathway separation results for three sets of related pathways: type I and type II interferon (IFN), neutrophil- and monocyte-/macrophage-related gene sets (MYELOID) and the G1 and G2 phases of the cell cycle (proliferation). Models were trained on either 5 randomly selected expression matrices of the same size (Sample Size) or on specific biological conditions with different random seeds (Biological Context). The cells of the heatmap are colored based on the number of models where separation for the related pathways is achieved or the presence or absence of separation in the MultiPLIER model. (There is only one MultiPLIER model.)

In the random subsampling experiment, a larger sample size resulted in a larger number of models achieving pathway separation (Figure 5). Separating the G1 and G2 phases of the cell cycle appeared to be particularly sensitive to the training set and starting conditions (Figure 5). Pathway separation was not just a function of training set sample size, as blood and tissue models perform well at IFN and myeloid separation, whereas the cell line training set was ill-suited for pathway separation for all three sets. MultiPLIER successfully separates the IFN, myeloid, and proliferation pathways (Figure 5).

We observed that the MultiPLIER model distinguished two types of IFN signaling (Figures 5 and S4A), which can be difficult to parse at the gene expression level (Chiche et al., 2014), whereas the SLE WB PLIER model (n = 1640) only learned a latent variable associated with type I IFN (Figure 2A). The SLE WB compendium includes two clinical trials of targeted treatments that block IFN signaling. We found that the MultiPLIER type I IFN-related latent variable (Figure S4A; note that the p-values displayed in Figure S4 are uncorrected) stratified patients into groups consistent with the original publication of an agent that blocks type I IFN signaling (Lauwerys et al., 2013). The MultiPLIER results were similar to a *whole blood-specific* modular transcriptional repertoire analysis approach popular in rheumatology studies of whole blood expression data (Figure S4B) (Chaussabel et al., 2008; Chiche et al., 2014; Oswald et al., 2015), whereas the SLE WB latent variable most associated with type I IFN signaling showed less evidence for change with treatment. The signatures (e.g., latent variables or modules) that captured type II IFN signaling showed a trend of decreased expression during treatment with a treatment that modulates this pathway (Welcher et al., 2015.), whereas the type I IFN signatures were unchanged (Figure S5).

### The latent space of the MultiPLIER model showed agreement with a PLIER model trained on a small vasculitis cohort and captures an autoantigen signature

The MultiPLIER approach is most attractive for rare disease where sample sizes are limited. AAV is a group of rare diseases characterized by neutrophil-mediated inflammation of small and medium vessels in multiple systems, and includes the sub-types of granulomatosis with polyangiitis (GPA), microscopic polyangiitis (MPA), and eosinophilic granulomatosis with polyangiitis (EGPA; formerly Churg-Strauss). We projected microarray datasets from multiple tissues in AAV—nasal brushings, microdissected glomeruli from kidney, and blood—into the MultiPLIER latent space and examined the expression of the MultiPLIER latent variables in these data. To our knowledge, the recount2 compendium contains no vasculitis data.

We evaluated the extent to which the MultiPLIER model, trained without any vasculitis data, captured vasculitis-relevant features. We trained a PLIER model on the NARES nasal brushing dataset (n = 79; Grayson et al., 2015), and compared the results to the projection into the MultiPLIER latent space. NARES includes data from patients with GPA, a form of AAV. First, we identified the MultiPLIER latent variable that was the best match for each of the 34 latent variables learned by the NARES PLIER as determined by correlation between the loadings (highest Pearson correlation coefficient). We observed that the expression levels of the best matched latent variables were positively correlated between the NARES and MultiPLIER latent spaces (i.e., *B* matrices) and much more so than we would expect than any random combination of latent variables between models (Figure 6). Figure S6 displays the correlation between latent variable expression alongside the correlation between loadings for the best match latent variables. Those NARES latent variables that are significantly associated (FDR < 0.05) with at least one gene set have particularly strong agreement (blue points in Figure 6).

**Figure 6.**
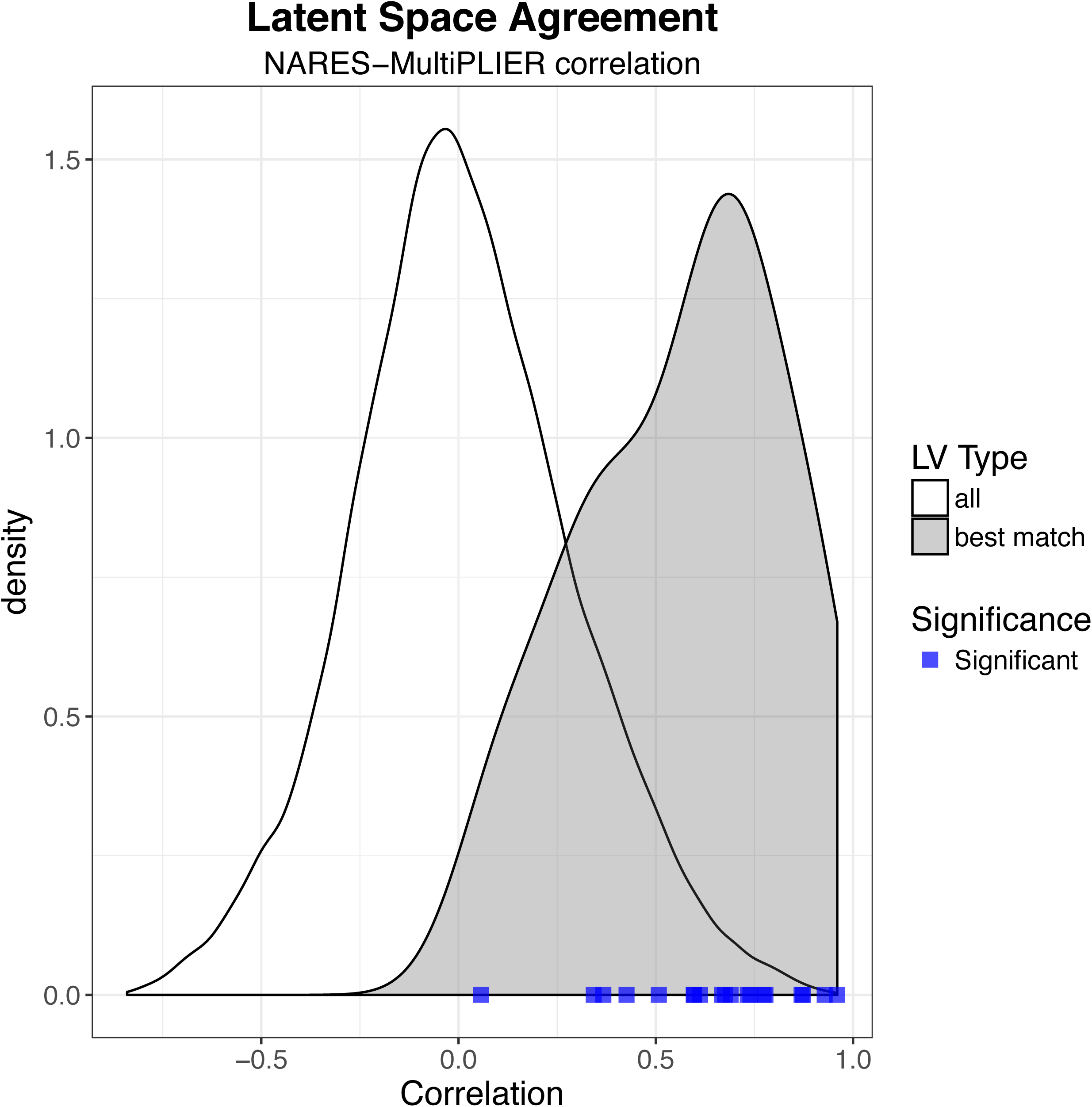
The latent space of the MultiPLIER model (no vasculitis data in the training set) shows agreement with the PLIER model trained on the NARES nasal brushing dataset (n = 79) particularly in the case of latent variables of biological significance. The best match for a NARES model latent variable was determined by identifying the latent variable (LV) in the MultiPLIER model with the most similar loadings (Pearson correlation of *Z*). NARES data was projected into the MultiPLIER latent space (*B*) and the Pearson correlation between *B* matrices (NARES, MultiPLIER) was calculated. Density plot of the latent variable expression value correlation coefficients between best match latent variables, shown in gray, demonstrates a rightward shift from the values for all pairs of latent variables between models (e.g., including random pairs of latent variables), shown in white. The blue points along the bottom of the graph represent best match correlation values for latent variables from the NARES model that are significantly associated (FDR < 0.05) with at least one gene set. This suggests that the values between latent variables that capture biological signal are particularly likely to be preserved in the transfer learning (MultiPLIER) case. Figure S6 shows the correlation values between best match latent variables alongside the correlation between loadings.

We identified differentially expressed MultiPLIER latent variables in the peripheral blood mononuclear cell (PBMC) fraction from Cheadle, et al. (Cheadle et al., 2010). This dataset included patients with GPA (known as Wegener’s granulomatosis at the time) and healthy controls. Patients with GPA were stratified into two groups in the original publication: those with expression patterns distinct from healthy control (here, termed “GPA-positive”) and those that were more similar to healthy controls (termed “GPA-negative”). We tested for differential expression between these three groups—GPA-positive, GPA-negative, and control—and identified MultiPLIER LV599 as differentially expressed (FDR = 1.17e-08; Figure S7A). ANCA antigens (i.e., the neutrophilic granule components that autoantibodies react with) are among the genes that contribute most to MultiPLIER LV599 (black bars in Figure S7D).

### A MultiPLIER meta-analysis framework identified differentially expressed latent variables in three tissues affected by ANCA-associated vasculitis

We were interested in molecular processes perturbed in the systemic disease AAV. Because we projected the three tissue datasets into the same latent space, MultiPLIER lends itself naturally to the meta-analysis of the NARES, glomeruli, and blood data as we did not have to attempt to match latent variables across models making analysis much less time consuming for multiple datasets. PLIER reduces technical noise (Figure 2C) and discards genes and gene sets that are irrelevant to the training data (Mao et al., 2017), which are distinct advantages over other gene set methods such as Gene Set Enrichment Analysis (GSEA; Subramanian et al., 2005). We performed differential expression analysis to identify differentially expressed latent variables in each of the three datasets and to find those latent variables that had similar expression patterns across tissues.

We examined signatures of more severe or active disease in AAV across organ systems. In the blood dataset, patients with a GPA-positive signature (most dissimilar from healthy controls) had higher Birmingham Vasculitis Activity Scores (BVAS; Cheadle et al., 2010). In the NARES dataset, there were three groups of patients with GPA—those with *active* nasal disease, those with *prior* nasal disease, and those that had *no history* of nasal disease—and a composite comparator group of patients with eosinophilic granulomatosis with polyangiitis (EGPA, Churg-Strauss), allergic rhinitis, sarcoidosis, and healthy controls (Grayson et al., 2015). The glomeruli dataset contained patients with AAV as well as those with nephrotic syndrome and living donor controls (Grayson et al., 2018). We used all available groups for comparisons (see STAR Methods). We identified 22 MultiPLIER latent variables that are differentially expressed (FDR < 0.05) in all three datasets and additional latent variables differentially expressed in two out of three tissues (Figure S7A).

We were particularly interested in latent variables that showed evidence of increased expression in GPA with active disease (NARES/nasal brushings), the GPA-positive group (PBMCs), and in the vasculitis group in kidney. MultiPLIER LV10, which is significantly associated with the M0 macrophage gene set from Newman, et al. (Newman et al., 2015), was overexpressed in these three groups relative to controls (Figure 7A-C; multigroup comparison FDR < 0.05; p-value from Wilcoxon rank sum test shown in figure). Top (highest weight) genes for MultiPLIER LV10 (Figure 7B) include the M2 macrophage marker *CD163* and *CCL2* which has been shown to influence macrophage polarization to an M2 phenotype (Sierra-Filardi et al., 2014). The top 25 genes also include genes related to tissue remodeling such as *MMP9* and *MMP14* (Figure 7D). MultiPLIER LV937 was significantly associated with a HIF-1a transcription factor network and showed a differential expression similar to MultiPLIER LV10 pattern (Figure S8B-D).

**Figure 7.**
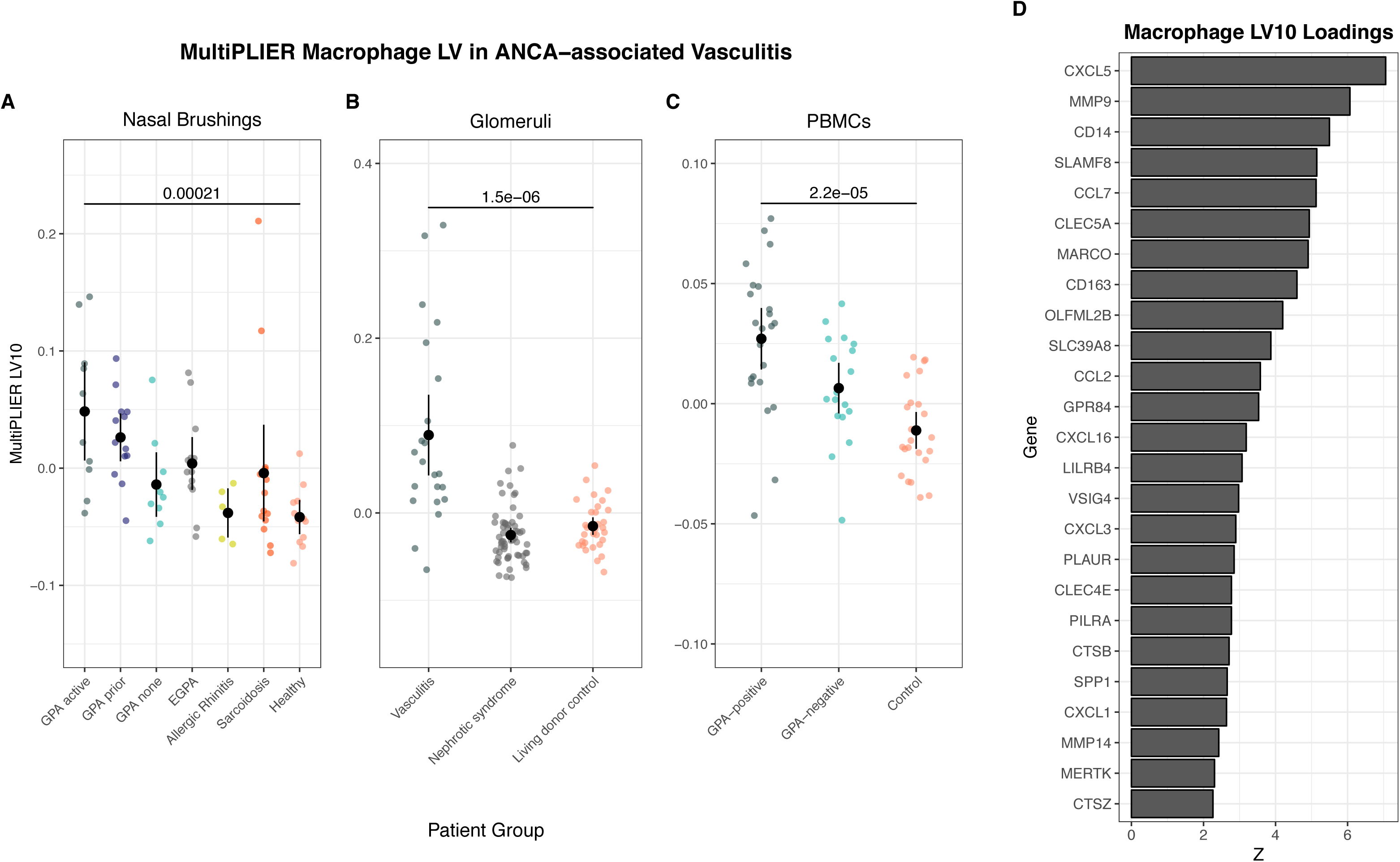
A MultiPLIER-learned latent variable associated with macrophages is differentially expressed in three tissues from ANCA-associated vasculitis (AAV) and shows increased expression in severe or active disease. Differentially expressed latent variables (LVs) were identified by comparing all patient groups and using Benjamini-Hochberg correction (FDR). Latent variables with FDR < 0.05 were considered to be differentially expressed. MultiPLIER LV10 had an FDR < 0.05 in all three cohorts. **(A-C)** Jitter plots of MultiPLIER LV10 in three different tissues: nasal brushings (NARES dataset), kidney microdissected glomeruli, and peripheral blood mononuclear cells. Points and bars represent mean ± 2 * SEM. P-values are from a Wilcoxon rank sum test comparing the control group to the AAV group considered to have the most severe or active disease in the cohort. **(D)** The loadings of the top (highest weight) 25 genes for MultiPLIER LV10.

### MultiPLIER latent variables were differentially expressed between medulloblastoma subgroups

The diseases examined thus far, SLE and AAV, have a strong autoimmune component and many of the gene sets supplied to the MultiPLIER model during training were related to immune cell sets. To evaluate how generalizable the MultiPLIER approach is to other types of rare disease, we analyzed two medulloblastoma data sets (Northcott et al., 2012; Robinson et al., 2012). To our knowledge and according to MetaSRA annotations, there were no medulloblastoma samples included in recount2.

Medulloblastoma can be classified into 4 subgroups on the basis of gene expression profiling: SHH, WNT, Group 3, and Group 4 (Taylor et al., 2011). The Northcott, et al. and Robinson, et al. data sets included subgroup labels (Northcott et al., 2012; Robinson et al., 2012). We tested for latent variables that were differentially expressed between subgroups. We found that 617 of 987 MultiPLIER latent variables were differentially expressed (FDR < 0.05) in both data sets (no guarantee of agreement with respect to directionality). Of the top differentially expressed latent variables that were associated with pathways in the Northcott, many were associated with processes related to translation. In Figure 8, we display two latent variables (MultiPLIER LV161 and LV707) associated with translation-related pathways that were differentially expressed in both cohorts, consistent with previous observations at the mRNA and protein level (Forget et al., 2018).

**Figure 8.**
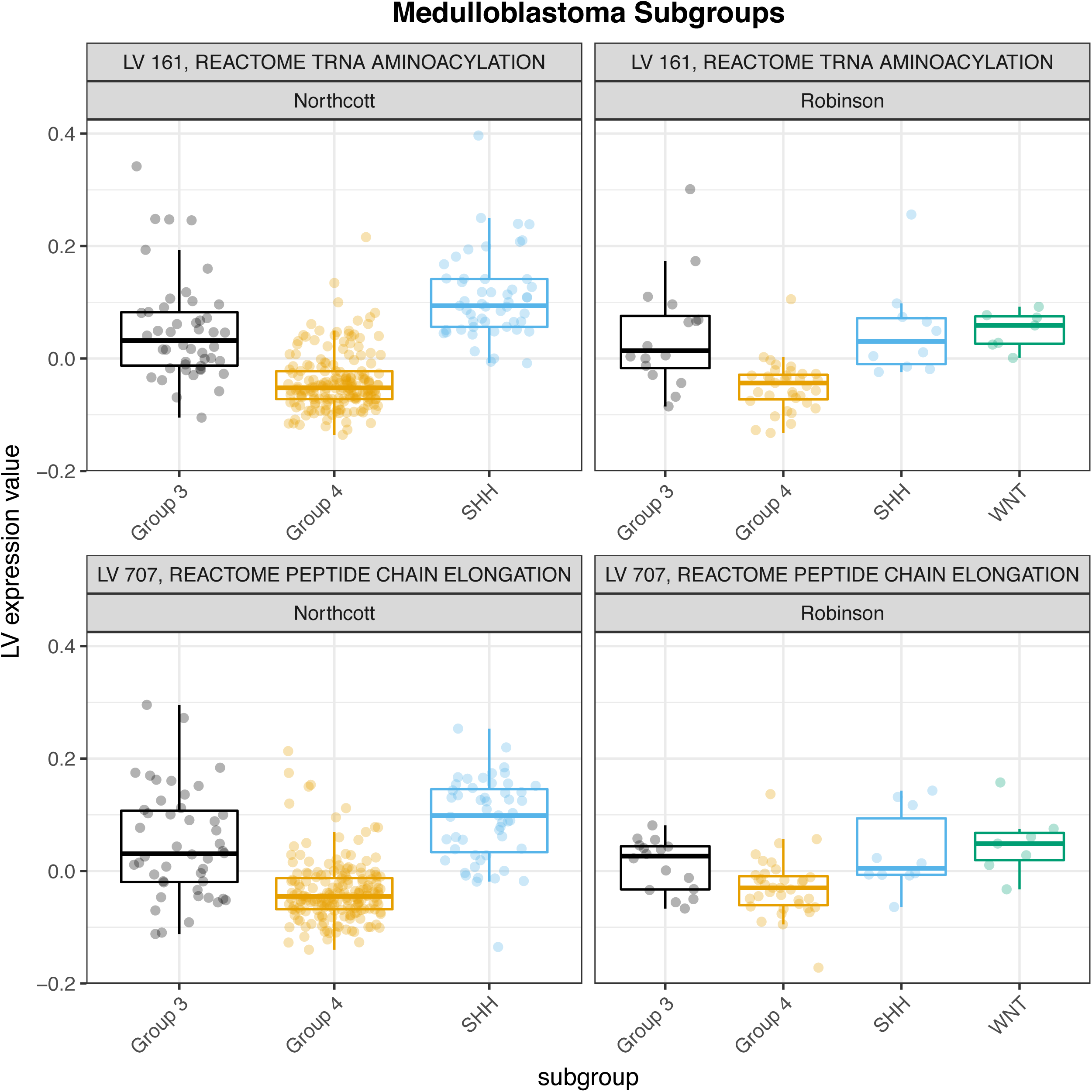
MultiPLIER latent variables associated with translation-related pathways are differentially expressed between subgroups in two medulloblastoma cohorts. Differentially expressed latent variables (LVs) were identified by comparing all patient groups and using Benjamini-Hochberg correction (FDR). Latent variables with FDR < 0.05 were considered to be differentially expressed. Expression data and subgroup labels from (Northcott et al., 2012) and (Robinson et al., 2012).

## DISCUSSION

We describe MultiPLIER—an unsupervised transfer learning framework for identifying perturbed molecular processes in complex human disease across organ systems. MultiPLIER models are generated by applying a robust factorization approach named PLIER (Mao et al., 2017) to a large, uniformly processed body of gene expression from multiple tissues and biological processes (Collado-Torres et al., 2017). We demonstrated that PLIER learned highly biologically relevant latent variables when trained on a single-disease, single tissue compendium comprised of multiple datasets on different microarray platforms. Our sample size and biological context experiments further support the use of our approach—namely, the use of multiple datasets, tissues, and biological conditions, to train a PLIER model with attractive properties. We demonstrated that models trained on larger sample sizes and on samples that we expect to contain a mixture of cell types had a higher number of latent variables and higher pathway coverage. Models trained on larger sample sizes and on particularly relevant biological contexts were better able to distinguish similar pathways.

We found that as sample size increased, a smaller proportion of latent variables were associated with pathways, with the exception of the blood training set (Figure 4C). The blood training set is well-suited to capture the input pathways because many of the pathways supplied to PLIER during training are immune cell-related and is likely to be the most homogeneous of the training sets with respect to the overall variety of cell types. The decrease in this proportion most likely cannot be explained by increased multiple testing burden alone (i.e., more latent variables, more hypotheses). If that were the case, we would expect the models trained on tissue samples to have lower values than those trained on cell line samples in Figure 4C. This suggests that more signal that is irrelevant with respect to the gene sets supplied during training is introduced when more samples and/or a greater breadth of samples are used as part of the training set. However, the MultiPLIER model is able to capture held out pathways, has high pathway coverage relative to other models, and is likely more applicable to a breadth of downstream applications given its large and diverse training set. For instance, models trained on the cancer data set did not NK cell signatures, which may be attributable to the lack of NK cell infiltrate in the included tumor samples and/or the proportion of cell line experiments in this training set.

MultiPLIER captured cell type-specific patterns as well as a model trained on the SLE whole blood compendium, despite the inclusion of very few SLE samples in recount2 and the cross-platform (RNA-seq training, microarray transfer) application, and was better at separating IFN pathways. The latent space results from the NARES AAV nasal brushing dataset agree strongly with the MultiPLIER latent space results, particularly in the case of pathway-associated latent variables. MultiPLIER captured biology shared in multiple tissue contexts (i.e., granule development) that is aberrant in GPA (autoantigens) without being trained on GPA data. The MultiPLIER model learned latent variables that were significantly associated with macrophage and HIF-1alpha TF network gene sets and had increased expression in more severe or active AAV in multiple tissues. This is consistent with the results of Grayson, et al., where we demonstrated signature associated with aberrant metabolism is present in AAV glomeruli and identified macrophages as the most likely cellular source (Grayson et al., 2018). Although we highlight these two latent variables, additional patterns of interest were identified (Figure S8A). Analysis of two medulloblastoma cohorts identified translation-related processes as differentially expressed between subgroups. Thus, we have demonstrated that training a model on data that represents an arbitrary sampling of human physiological processes captures well-documented features similarly to a tissue- and disease-specific model in a common disease, and learns additional biological factors or pathways, which aids in the interpretation of rare disease expression data from multiple tissues.

There are several limitations of this study. For instance, we made minimal effort to adjust the data distributions between platforms (i.e., RNA-seq and microarray) (Tan et al., 2017; 2016; Thompson et al., 2016) or impute the expression of genes missing between models. Despite these areas for improvement, studying the expression of MultiPLIER latent variables in microarray datasets from rare disease proved to be useful. We also presented the results for one MultiPLIER model trained on the recount2 compendium and PLIER is non-deterministic. An ensemble approach, like that introduced in Tan, et al. (Tan et al., 2017) could be beneficial, though our repeats of biological context PLIER training suggest that PLIER produced stable models at least in terms of the number of latent variables and proportion of input pathways captured. Additionally, any approach that transfers patterns learned in a large compendium to an “unseen” disease dataset runs the risk of missing subtle disease-specific and disease-affected tissue-specific signals. Conducting hypothesis-generating research in small datasets in general poses a risk for false discovery. After first comparing model features to well-characterized datasets to validate model construction, we focus on patterns that are present in multiple datasets and tissues.

PLIER leverages the underlying modular structure of gene expression data. MultiPLIER expands on this by taking advantage of the biological processes shared among different conditions. We expect that the benefits of training on a large compendium of uniformly processed data—the first of which is the reduction of human curation and processing time associated with assembling a disease-specific compendium even when feasible—are not limited to PLIER. We speculate that other matrix factorization approaches and autoencoders that have been shown to extract a biologically meaningful low dimensional representation from gene expression data (Dincer et al., 2018; Stein-O’Brien et al., 2017b; Tan et al., 2017; Way and Greene, 2017) are appropriate for a similar transfer learning approach. The constraints that force the PLIER model to learn biological signal made it an attractive method for this work. Indeed, the latent variables learned by MultiPLIER that were significantly associated with gene sets were shown to be in strong agreement with dataset-specific model latent variables. Future investigation into a variety models with different strengths (e.g., those that capture nonlinear patterns), with particular emphasis on the factors in the latent space that are well-aligned with biological signatures of interest, is supported by this work.

Although we propose the MultiPLIER is a particularly strong tool for the study of rare diseases, in which the size of gene expression datasets are inherently small compared to similar data derived from common diseases, we also foresee that MultiPLIER might be quite useful for the increasing study of precision (personalized) medicine and the now routine recognition of newly-discovered single-gene subsets of diseases. The precision medicine approach is linked to the growing appreciation of the importance of identifying subsets of patients within larger populations of patients with a specific disease. Such subsets may identify patients with i) different sets of clinical manifestations of disease; ii) varying degrees of disease severity; iii) different long-term outcomes; and iv) varied responses to specific therapies. Such disease subsets almost certainly have some differences in genetic background and/or gene expression in involved tissues. Thus, these approaches to precision medicine will essentially transform some common diseases into subsets of rarer diseases, many of which could benefit from the MultiPLIER approach to data analysis.

## METHODS

### Gene expression data

#### recount2 compendium

We obtained gene-level RangedSummarizedExperiment data from the recount2 compendium (Collado-Torres et al., 2017) using the recount Bioconductor package and calculated Reads Per Kilobase Million (RPKM). We excluded samples without metadata; a total of 37,032 samples were included. Individual experiments were concatenated without any additional processing and genes were z-scored for use with PLIER. We converted from Ensembl gene IDs to gene symbols using the biomaRt Bioconductor package.

To describe the subset of recount2 data that was used to train the model, we used MetaSRA database (Bernstein et al., 2017) (v1.4). MetaSRA uses a custom pipeline to predict sample source (e.g., cell line) and label with ontology terms from sources such as the Experimental Factor Ontology (Malone et al., 2010). We mapped run accessions (utilized in recount2) to sample accessions (utilized in MetaSRA) with the SRAdb (Zhu et al., 2013). Approximately 99% of samples in the training compendium were also present in MetaSRA. We describe the terms used for the biological context experiments in Table 2.

**Table 2.** Terms used to subset recount2 into training sets for the biological context experiments and the set sample sizes. Labels are from MetaSRA (Bernstein et al., 2017).

| Training set | Term or strategy used | Sample size (n) |
| --- | --- | --- |
| Blood | UBERON:0000178 | 3862 |
| Cancer | EFO:0000311 | 8807 |
| Tissue | MetaSRA sample type = tissue | 12396 |
| Cell line | MetaSRA sample type = cell line | 14532 |
| Other tissues | Everything but samples that mapped to blood training set | 32984 |

#### Microarray data

The NARES nasal brushing dataset, the isolated immune cell data (E-MTAB-2452) and the following systemic lupus erythematosus datasets were normalized using Single Channel Array Normalization (SCANfast implementation) and Brainarray v22.0.0 (Entrez ID version; Dai, 2005): E-GEOD-39088, E-GEOD-61635, E-GEOD-72747, and E-GEOD-11907. The medulloblastoma datasets (GSE37382, GSE37418) were processed with SCANfast and Brainarray v22.0.0 (Ensembl gene ID version; Dai, 2005) through refine.bio (www.refine.bio) without quantile normalization and without any gene-wise transformation; we selected all samples from the experiment that were available at the time of analysis. The NARES dataset was adjusted for batch effect with ComBat (Johnson et al., 2007). The glomeruli dataset was annotated using Brainarray v19.0.0, quantile normalized, and batch corrected using ComBat as in (Grayson et al., 2018). The publicly available submitter-processed data was used for all other datasets (e.g., SLE WB [E-GEOD-49454, E-GEOD-65391, E-GEOD-78193], GPA PBMCs [GSE18885]). The GPA PBMCs data was z-scored for use with PLIER and MultiPLIER. All conversions from Entrez or Ensembl gene IDs to gene symbols (the input to PLIER) were performed using the org.Hs.eg.db Bioconductor package.

#### Systemic lupus erythematosus whole blood compendium

The Illumina and Agilent submitter-processed data gene identifiers were converted to Entrez IDs using the illuminaHumanv4.db and hgu4112a.db packages, respectively. Duplicate Entrez IDs were collapsed to the mean expression value. Paired samples from E-GEOD-11907 (Affymetrix HGU133A and HGU133B) were concatenated. All SCAN-processed Affymetrix data was concatenated and the submitter-processed datasets were normalized using the Affymetrix target quantiles (preProcessCore Bioconductor package). Genes were [0,1] scaled in each dataset before joining; only genes present in each dataset were retained.

### Training Pathway-Level Information Extractor (PLIER) models and evaluation

#### PLIER training

We used the PLIER R package (commit: a2d4a2) to train PLIER models. The number of latent variables (*k* parameter) was set by determining the number of significant principal component using the PLIER num.pcs function and, following the authors recommendations in a version of the package documentation (https://github.com/wgmao/PLIER/blob/125407dd68fc95c5b771b65ae2eb237b629c4e89/vignett es/vignette.pdf), included an additional 30%. The expression matrix used as input to the PLIER function was z-scored using the PLIER rowNorm function. We used the bloodCellMarkersIRISDMAP, svmMarkers, and canonicalPathways data included in the package as the prior information. We used the default settings of the PLIER function, which automatically sets the L1 and L2 parameters (see Mao et al., 2017 for more information about implementation).

#### Random subsampling and biological context experiments

For the sample size experiments (Figure 4), we randomly selected 500, 1000, 2000, 4000, 8000, 16000, and 32000 samples from the recount2 dataset as our new expression matrix and treated it as described above. We trained 5 PLIER models on the subsampled recount2 dataset and each of the biological context training sets (blood, cancer, tissue, cell line, other tissues) initialized with different random seeds. We used the sample type labels, Uberon (Mungall et al., 2012)or Experimental Factor Ontology (Malone et al., 2010) terms in MetaSRA (Bernstein et al., 2017) to subset recount2 into the biological context training sets (Table 2). Training was performed as described above. To evaluate the models, we calculated the proportion of input pathways that were significantly associated (FDR < 0.05) with at least one latent variable—termed “pathway coverage”—and the proportion of the total latent variables that were significantly associated (FDR < 0.05) with at least one gene set.

When comparing what pathways or gene sets were captured by models trained on different biological contexts (e.g., blood vs. all other contexts), we considered a pathway/gene set to be captured by a biological context when 3 or more of the 5 models had a latent variable significantly associated (FDR < 0.05) with that pathway/geneset.

#### Pathway separation

We considered two sets of similar or related pathways/gene sets to be successfully separated by a PLIER model if the following conditions were met: 1) for each pathway set, at least one latent variable is significantly associated (FDR < 0.05) with a pathway in that set and 2) each pathway set is uniquely and significantly associated with at least one latent variable. Table 3 lists the gene sets used for this evaluation.

**Table 3.** Gene sets used for pathway separation evaluations.

| Evaluation | Set 1 | Set 1 gene sets | Set 2 | Set 2 gene sets |
| --- | --- | --- | --- | --- |
| Interferon (IFN) | Type I IFN | REACTOME Interferon Alpha/Beta Signaling | Type II IFN | REACTOME Interferon Gamma Signaling |
| Myeloid | Neutrophil | DMAP GRAN3, IRIS Neutrophil-Resting, SVM Neutrophils | Monocyte/macrophage | IRIS Monocyte-Day0, IRIS Monocyte-Day1, IRIS Monocyte-Day7, SVM Monocytes, SVM Macrophages M0, SVM Macrophages M1, SVM Macrophages M2 |
| Proliferation | G1 phase | REACTOME G1 S Transition, REACTOME M G1 Transition, REACTOME, APC C CDH1 Mediated Degradation of CDC20 and other APC C CDH1 Targeted Proteins in Late Mitosis Early G1, REACTOME Cyclin E Associated Events During G1 S Transition, REACTOME G1 Phase, REACTOME Mitotic M M G1 Phases, REACTOME P53 Dependent G1 DNA Damage Response, REACTOME G1 G1 S Phases, Reactome P53 Independent G1 S DNA Damage Checkpoint | G2 phase | REACTOME Mitotic G2 G2 M Phases, REACTOME G2 M Checkpoints |

#### Held out pathway coverage

To determine MultiPLIER’s pathway coverage of the MSigDB oncogenic pathways (the oncogenicPathways data included in the PLIER package) (Subramanian et al., 2005), we used the same methodology as the PLIER crossVal and AUC (internal) functions to calculate AUC, p-values, and FDR values for pathway-latent variable associations; all pathway-latent variable associations were tested. We calculated pathway coverage as described above.

### MultiPLIER

We summarize the relevant details from the PLIER method below. No alterations to the underlying PLIER framework were made. For more information about implementation, see Mao, et al. (Mao et al., 2017). Given the gene expression matrix *Y* and the prior information matrix (e.g., gene sets) *C*, PLIER finds the prior information coefficient matrix *U*, the loadings matrix *Z*, and the latent variables matrix *B*, minimizing:

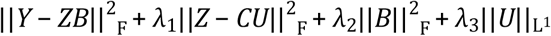

where *U >* 0 and *Z >* 0. The terms are minimizing the reconstruction error, enforcing that *Z* represents sparse combinations of the input gene sets, an L2 penalty on *B* and an L1 penalty on *U*, respectively. As noted above, we used the default settings for *λ*_1_ and *λ*_2_ for a given training set when training a PLIER model (see Mao et al., 2017 for more information). In practice, setting *λ*_2_ = 0 during training causes *Z* to no longer be sparse and a marked decrease in pathway coverage (see the notebook 39-L2_penalty in our Github repository for the full details and results of this experiment).

The final *B* matrix returned by PLIER is calculated as follows:

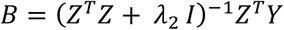

We use the *Z* matrix and the *λ*_2_ parameter from the source domain model to calculate the expression values of the target samples in the source domain latent space *B*_*target*_ below:

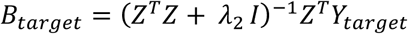

Where *Y*_*target*_ is the row-normalized (z-scored) expression matrix from the dataset (target domain) that we would like to project into the latent space learned by the source domain model. If genes present in the training set (source domain) were absent from the target gene expression matrix, we set the expression values for the missing genes to zero for all samples. As the expression data is z-scored, this is in essence setting the expression values to the mean. We calculated *B*_*target*_ for the following microarray datasets: SLE WB compendium, NARES nasal brushing, microdissected glomeruli (kidney), GPA peripheral blood mononuclear cells, medulloblastoma tumor samples, and isolated leukocytes (CD4, CD14, CD16 selection).

#### Latent variable differential expression analysis

We performed differential expression analysis using the limma Bioconductor package. We performed contrasts between all groups and used Benjamini-Hochberg adjustment for multiple hypothesis testing (“BH” method). We considered all latent variables with FDR < 0.05 to be differentially expressed.

### Principal components analysis

We performed principal components analysis (PCA) on the fully processed SLE WB compendium (Figure S1), as well as subsets of the latent space (*B* matrix) from the SLE WB PLIER model using the prcomp function in R. The subsets of *B* were as follows: all latent variables (“All Latent Variables” Figure 2B), only latent variables with AUC > 0.75 for at least one gene sets (“Pathway-associated Latent Variables” Figure 2C), only latent variables with AUC ≤ 0.75 for all gene sets (Figure 2D). We repeat this with different AUC values in Figure S1.

### Cell type evaluations

Neutrophil-associated latent variables were identified based on the AUC and FDR (< 0.05) values with neutrophil gene sets. We selected neutrophil-associated latent variables for display in Figure 3 based on a lack of significant association with monocyte-, macrophage-, or granulocyte progenitor-related gene sets, as this suggests specificity for neutrophils rather than a general myeloid lineage association. We used the neutrophil counts from the metadata associated with the publicly available accession (E-GEOD-65391) for Banchereau, et al. (Banchereau et al., 2016). We calculated the MCPcounter neutrophil estimate with the MCPcounter.estimate function in the MCPcounter R package (Becht et al., 2016). We used the R linear model function (lm) to calculate the *R*^2^ values in the neutrophil scatterplots in Figure 3. Disease activity group labels (Figure S2) were also obtained from the E-GEOD-65391 sample metadata. Plasma cell-associated latent variables were identified by their significant (FDR < 0.05) association with plasma cell gene sets; latent variables with the highest AUC were selected for display. Pairwise t-tests between disease activity groups were performed with the pairwise.t.test function in R. For the heatmap in Figure S3, we selected latent variables for display if they had one of the following strings in their names: CD4, monocyte, or neutrophil.

### Interferon clinical trials

#### Whole blood modular framework

We used the IFN-related modules reported in Chiche, et al. (Chaussabel et al., 2008; Chiche et al., 2014; Obermoser et al., 2013). Briefly, in this approach, modules are constructed by clustering gene expression data from whole blood from patients with a variety of autoimmune or infectious conditions and identifying shared expression patterns (Chaussabel et al., 2008). The result is sets of genes with correlated expression or modules. It is important to note that these modules are then somewhat specific to whole blood. We obtained the gene sets from http://www.biir.net/public_wikis/module_annotation/V2_Trial_8_Modules and converted to Entrez ID using Tribe (Zelaya et al., 2016). (These converted gene sets are available in our analysis Github repository.) We summarized the expression of a module by taking the mean expression value for genes in the module for a sample.

#### PLIER models

We selected IFN-related latent variables based on their association with the following genesets: REACTOME Interferon signaling, REACTOME Interferon alpha/beta signaling (Type I), REACTOME Interferon gamma signaling (Type II) (Fabregat et al., 2017; Subramanian et al., 2005).

#### IFN-K trial analyses

In our analyses of the IFN-alpha-kinoid trial (E-GEOD-39088; Lauwerys et al., 2013), we stratified the patients into IFN-positive and IFN-negative groups to be consistent with the original publication and therefore facilitate our evaluations. We did not have the labels from the original publication. We identified IFN-negative samples based on baseline expression values and using the following criteria: 9 patients with the lowest summary expression values for M1.2 (whole blood modular framework), 9 patients with the lowest expression of MultiPLIER LV116 (MultiPLIER), or 9 patients that had the lowest expression in at least 2 of the SLE WB latent variables—LV6, LV69, LV110 (SLE WB PLIER). All other patients were considered IFN-positive. Patients that received placebo were placed in a “placebo” group for analysis. We calculated the change in expression value between baseline and days 112 and 168 for each patient.

### Latent space agreement

To determine whether or not the NARES PLIER model and MultiPLIER latent space expression values were in agreement, we first identified the best match latent variables between the two models. For each pair of latent variables between the two models, we calculated the Pearson correlation coefficient between the loadings (*Z*). The MultiPLIER latent variable with the highest correlation value to an individual latent variable from the NARES model was considered the best match. We calculated the Pearson correlation values between the *B* matrix learned by the NARES PLIER model and *B* from the projection of the NARES data into the MultiPLIER latent space.

## ACKNOWLEDGMENTS

We would like to thank Gregory P. Way for code review and thoughtful discussion.

## DATA AND SOFTWARE AVAILABILITY

Code for expression data processing and MultiPLIER training is available at https://github.com/greenelab/rheum-plier-data and at the following accessions: E-GEOD-65391, E-GEOD-11907, E-GEOD-39088, E-GEOD-72747, E-GEOD-49454, E-GEOD-61635, E-GEOD-78193, GSE104948, GSE119136, GSE37382, and GSE37418. Code for analysis is available at https://github.com/greenelab/multi-plier. A release (v0.2.0) of the data, code, and models is available at DOI: 10.6084/m9.figshare.6982919.v2. We have prepared a Docker image with the software dependencies required to reproduce the analysis: https://hub.docker.com/r/jtaroni/multi-plier/ (tag 0.2.0).

## SUPPLEMENTAL FIGURE LEGENDS

**Figure S1.**
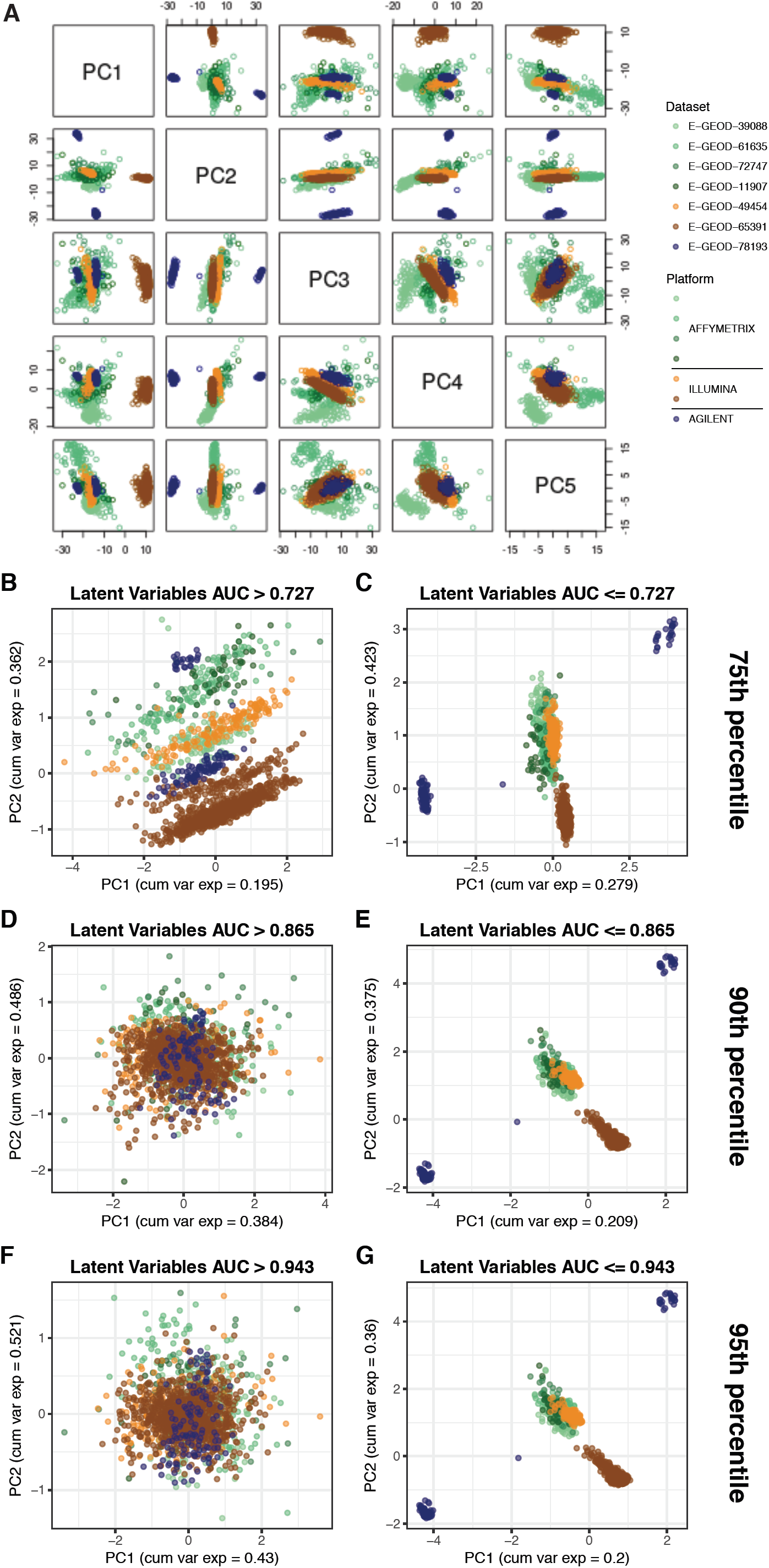
Principal components 1-5 for the SLE WB expression compendium. Related to Figure 2. Points are samples. Samples are colored by dataset of origin and datasets from the same platform manufacturer are similar colors. PC1-5 account for 0.502 of the variance explained. Panels B-G display the first two PCs from different subsets of the latent space or *B* matrix, either those associated with input gene sets (e.g., above the labeled AUC threshold) or those that are not. AUC thresholds were chosen based on the distribution of all AUC values.

**Figure S2.**
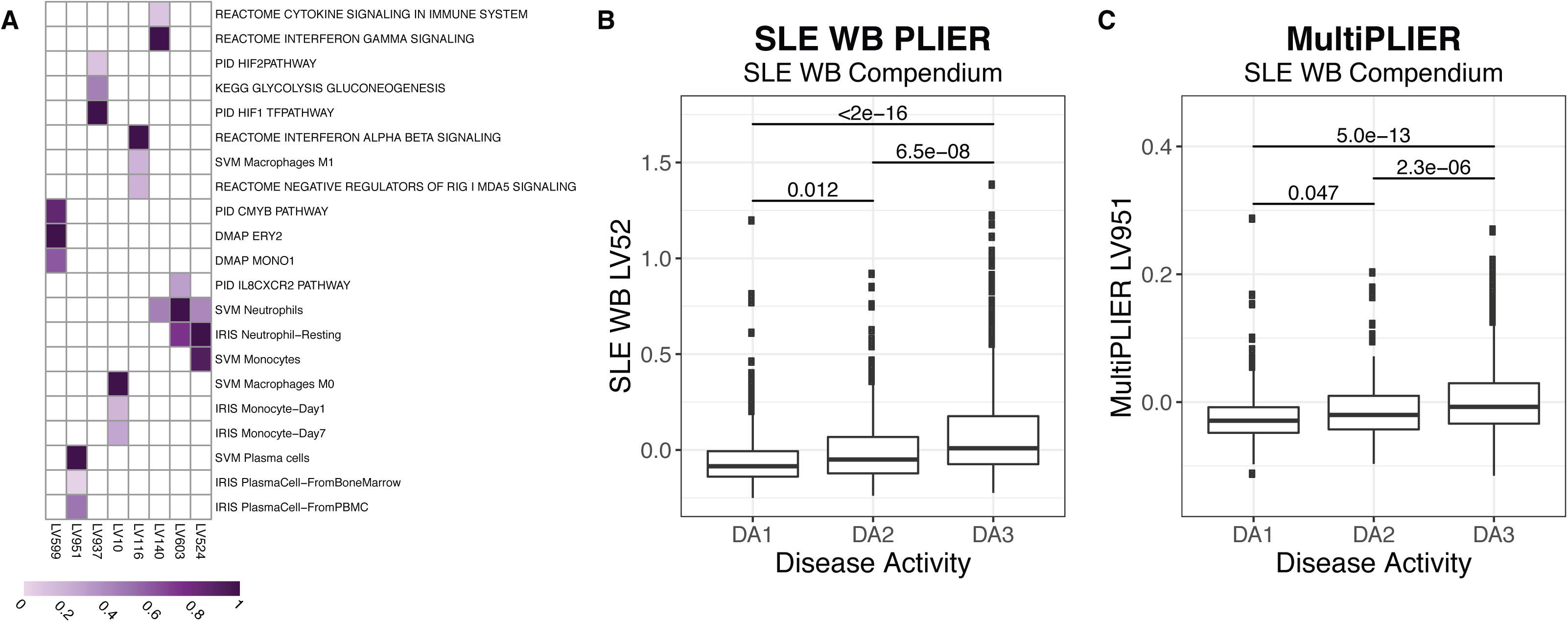
MultiPLIER learns a plasma cell signature that is elevated in more severe disease. Related to Figure 3. **(A)** Selected latent variables (LVs) from the MultiPLIER *U* matrix. Purple fill in a cell indicates a non-zero value and a darker purple indicates a higher value. Only pathways with FDR < 0.05 in displayed latent variables are shown. We selected latent variables discussed in the main text. **(B-C)** In Banchereau, et al. citation the authors stratified patients into three disease activity groups—DA1 (SLEDAI: 0-2), DA2 (SLEDAI: 3-7), and DA3 (SLEDAI > 7)— and demonstrated that patients with more severe disease had higher plasmablast counts (via FACS). We did not provide a plasmablast-specific gene set to the PLIER model during training. However, plasma cell from signatures adapted from the Immune Response In Silico (IRIS) project (Abbas et al., 2005) and Newman, et al. (Newman et al., 2015)(included in the PLIER R package). A plasmablast is the precursor to a plasma cell and we therefore assume that these cell types share gene expression patterns. Both the SLE WB **(B)** and MultiPLIER **(C)** models learn a latent variable associated with this signature (SLE WB LV52 and MultiPLIER LV951, respectively). P-values are from a pairwise t-test with Bonferroni correction. Both latent variables show a similar pattern to the original publication FACS plasmablast counts. However, there is evidence that the MultiPLIER latent variable may more specific to plasma cells, as SLE WB LV52 is also significantly associated (FDR < 0.05) with REACTOME Unfolded Protein Response and REACTOME Asparagine N-linked Glycosylation (Fabregat et al., 2017)(Figure 2A). MultiPLIER LV951 is only significantly associated with plasma cell gene sets (data not shown).

**Figure S3.**
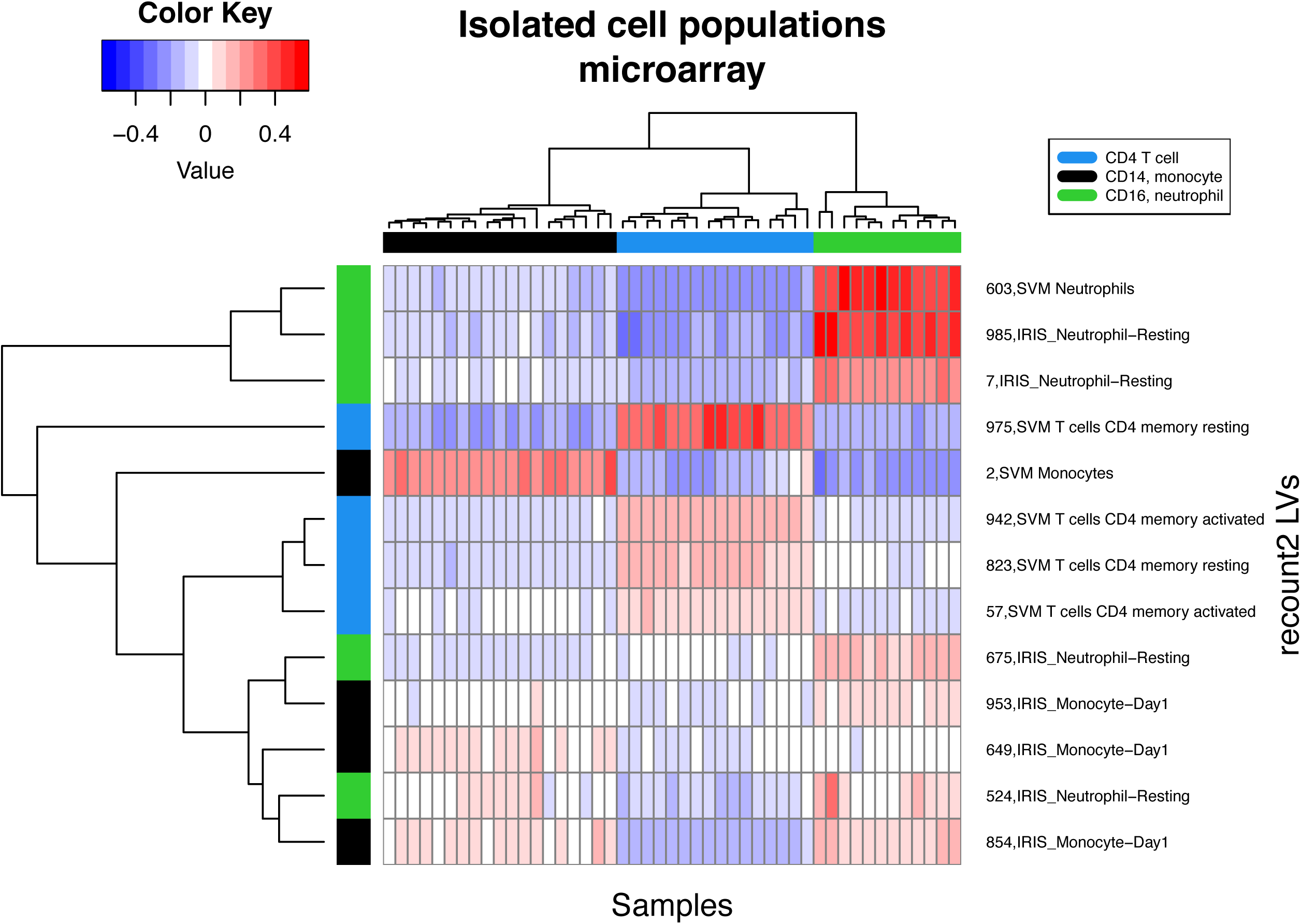
Projection of isolated leukocyte gene expression data, as measured by microarray, into MultiPLIER latent space demonstrates that the model captures additional cell type-specific signatures. Related to Figure 3. Heatmap of a subset of the B matrix for E-MTAB-2452 (McKinney et al., 2015), a dataset comprised of isolated immune subsets (CD4^+^ T cells, CD14^+^ monocytes, and CD16^+^ neutrophils) from patients with autoimmune diseases. Latent variables (rows) were selected if their names contained one of the following terms: CD4, monocyte, or neutrophil. The sample (column) annotation bar is colored based on the cell subset of that sample. The latent variable (row) annotation bar is colored by the term associated with the latent variable.

**Figure S4.**
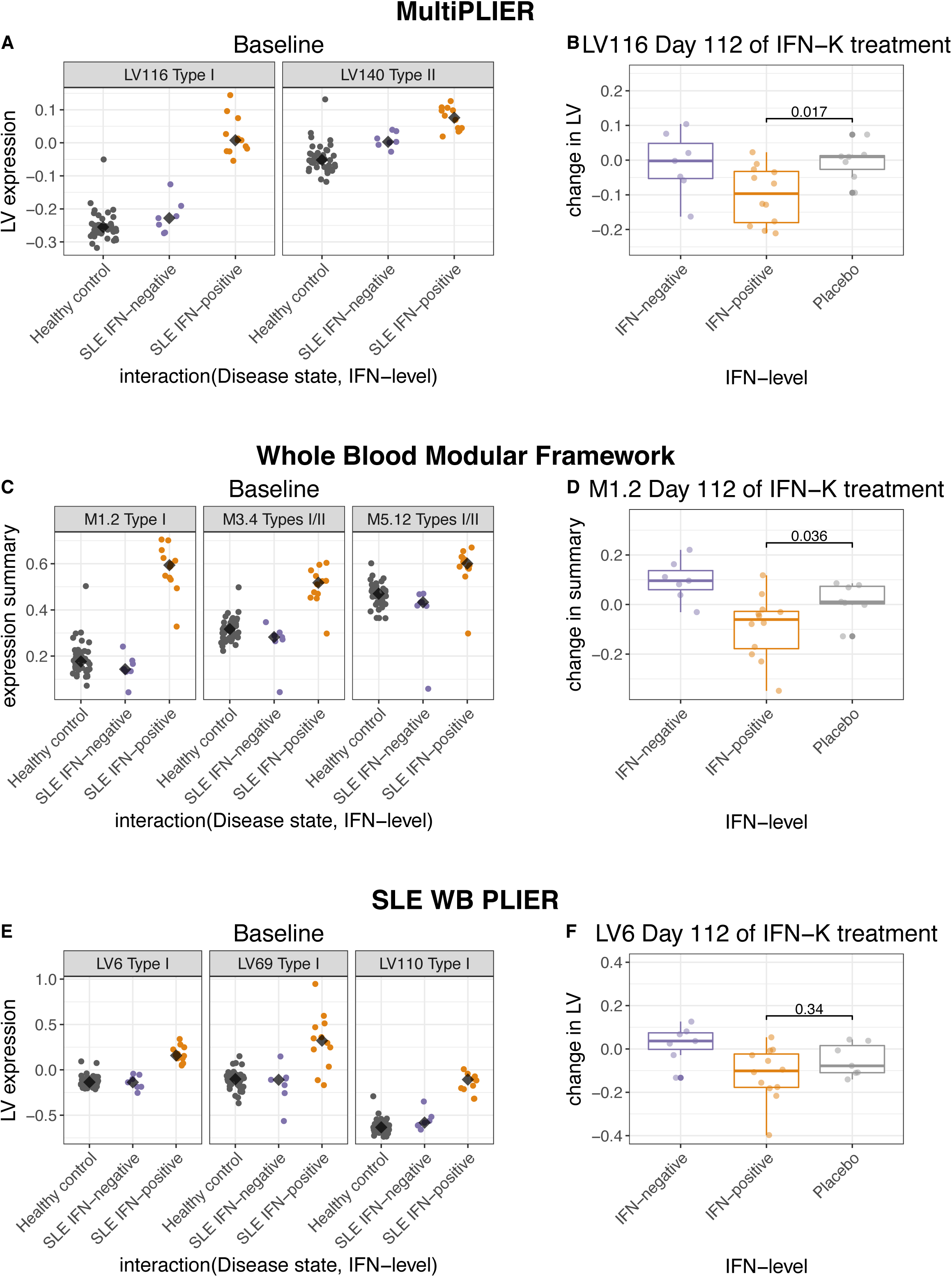
Expression patterns from IFN-K (type I IFN blockade) trial in the MultiPLIER latent space, whole blood modular framework, and SLE WB PLIER latent space. Related to Figure 5. Lauwerys, et al. (Lauwerys et al., 2013) is a clinical trial of the IFN-alpha-kinoid (IFN-K) therapeutic that results in the blockade of type I IFN via polyclonal antibody response. The authors of the original IFN-K publication stratified patients with SLE into two groups: IFN-negative (n = 9), who had IFN-alpha induced gene expression signatures that resembled healthy controls, and IFN-positive patients (n = 18), who had higher expression of the IFN-alpha signature they derived through *ex vivo* stimulation of healthy control blood with IFN-alpha. (Group labels from the original publication were not associated with the accession.) **(A)** Patients can be stratified into IFN-positive and IFN-negative groups using baseline expression of MultiPLIER LV116 (latent variable 116 from the MultiPLIER model). The expression of type II latent variable, MultiPLIER LV140, does not show a clear IFN-negative and IFN-positive group. **(B)** Treated IFN-positive patients show a reduction in MultiPLIER LV116 expression at day 112 as compared to the placebo group (p-value from Wilcoxon rank sum test). Chiche, et al. (Chiche et al., 2014) identified three IFN-related modules—M1.2, M3.4, and M5.12—and demonstrated that M1.2 was strongly induced by type I IFN, whereas the other two modules were similarly responsive to type I and type II IFN. **(C)** Type I module M1.2, and to a lesser extent M3.4, from the whole blood modular framework divides patients into IFN-positive and IFN-negative groups. Expression for each module was summarized using the mean expression value of all genes in the module. **(D)** Treated IFN-positive patients show a reduction in M1.2 expression at day 112 as compared to the placebo group (p-value from Wilcoxon rank sum test). **(E)** LV6, LV69, and LV110 from the SLE WB PLIER model were all significantly associated with type I IFN (REACTOME Interferon alpha/beta signaling). Patients were classified as IFN-negative if they had one of the nine lowest expression values in 2 out of 3 of these latent variables (typically LV6 and LV110). Recall that these samples are in the training data for this model. **(F)** Treated IFN-positive patients do not show a decrease in LV6 expression at day 112 as compared to placebo (p-value from Wilcoxon rank sum test). SLE WB LV6 was selected for display and comparison as it had the highest AUC for REACTOME Interferon alpha/beta signaling in the SLE WB model. Displayed p-values are not corrected for multiple hypothesis testing.

**Figure S5.**
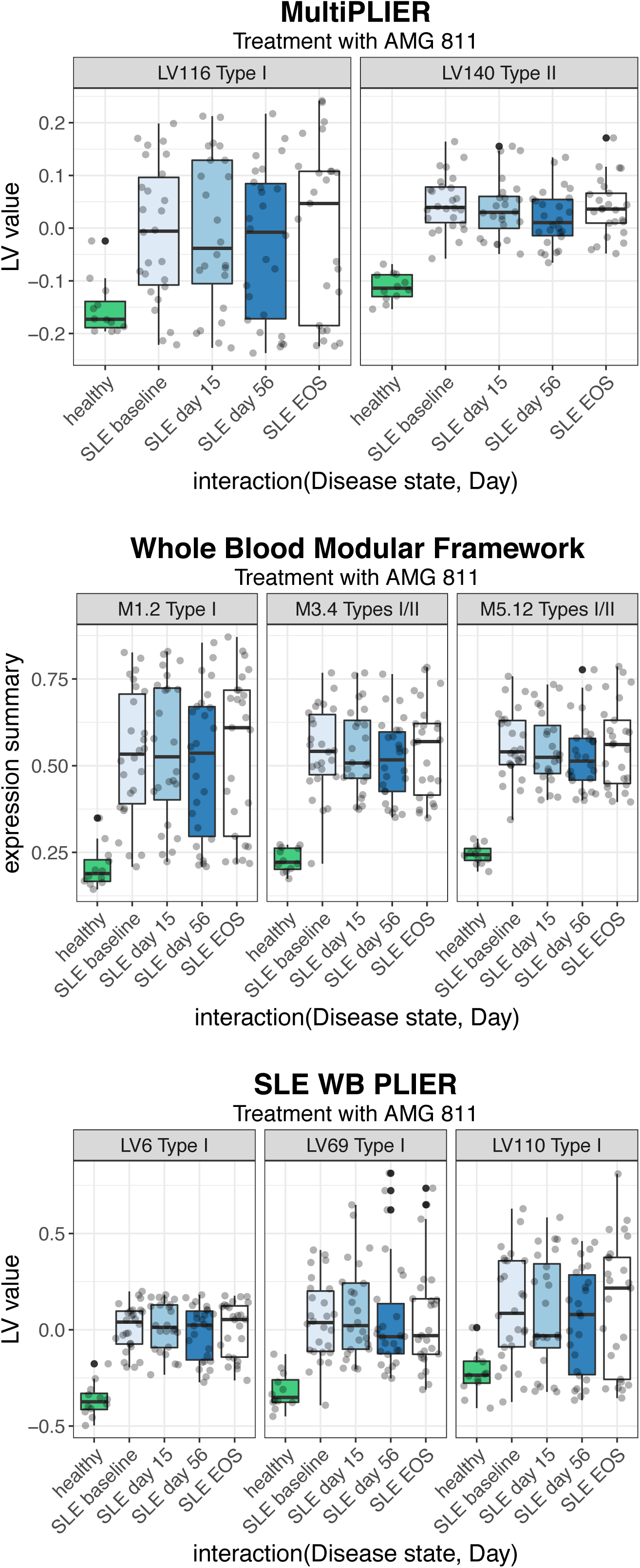
Expression patterns from trial of type II IFN blockade agent AMG 811 in the MultiPLIER latent space, whole blood modular framework, and SLE WB PLIER latent space. Related to Figure 5. The type II IFN latent variable (LV) from the MultiPLIER model, LV140, shows a trend towards decreased expression during days 15 and 56 of the AMG811 trial (top panel). The modules most likely to capture type II IFN expression, M3.4 and M5.12, show a similar trend (middle pattern). The latent variables or modules indicative of type I signaling show no such pattern (MultiPLIER LV116, whole blood modular framework M1.2, and all SLE WB latent variables). No SLE WB PLIER latent variable was significantly associated with type II (as determined by the lack of association with the REACTOME Interferon gamma signaling pathway).

**Figure S6.**
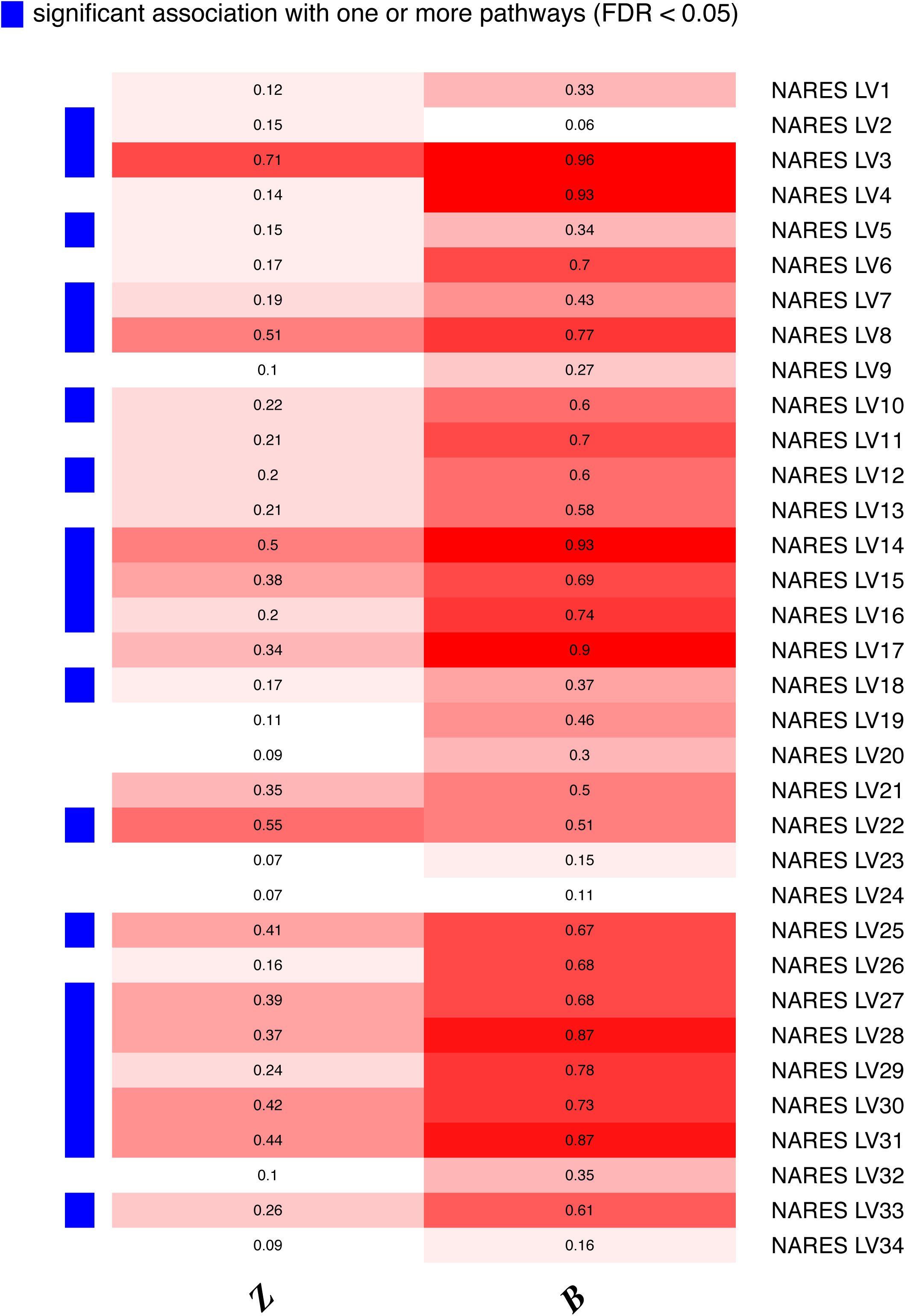
Comparison of the loadings (*Z*) and latent space (*B*) of the NARES PLIER model to the MultiPLIER loadings and the projection of the NARES dataset into the MultiPLIER latent space. Related to Figure 6. Heatmap of correlation values between the loadings and latent space of best match latent variables, where rows are the latent variables (LVs) from the NARES PLIER model. A more saturated red color indicates a higher correlation value. A NARES latent variable’s best match is determined by calculating the Pearson correlation between loadings (*Z* matrices) and selecting the latent variable from the MultiPLIER model with the highest correlation to that NARES latent variable. The columns are correlation values for the loadings (*Z*) and the latent variable expression (*B*) between that NARES latent variable (row) and its best match in the MultiPLIER model. The row annotation bar indicates whether or not a NARES latent variable is significantly associated (FDR < 0.05) with one or more gene sets supplied during training.

**Figure S7.**
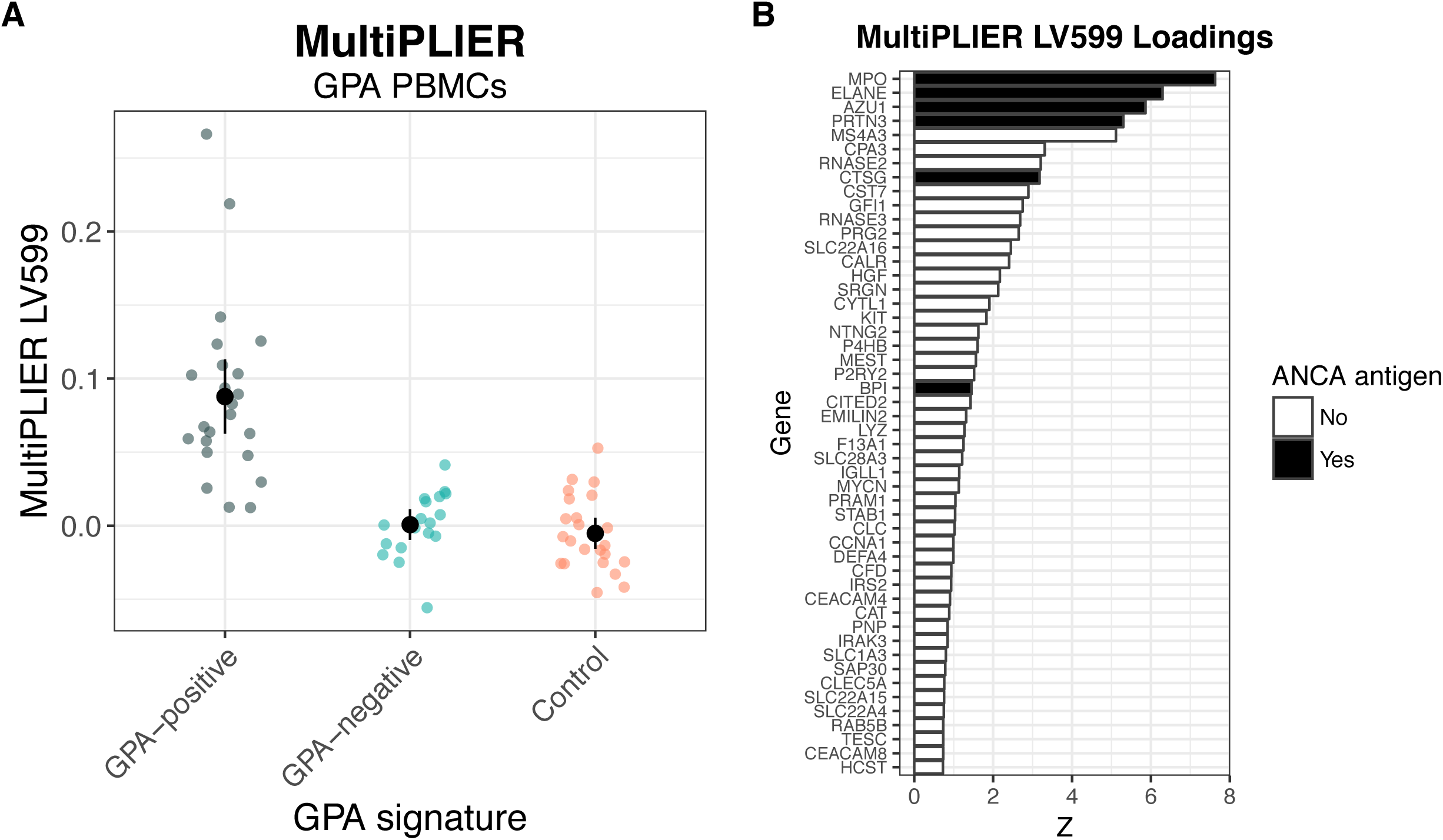
MultiPLIER learns a latent variable (LV) that captures the activity of antigens in antineutrophil cytoplasmic antibody (ANCA)-associated vasculitis despite no vasculitis training data. Related to Figure 7. The peripheral blood mononuclear cell (PBMC) fraction data from Cheadle, et al. (Cheadle et al., 2010) was projected into the MultiPLIER latent space. The authors of Cheadle, et al. identified three groups: GPA-positive (“WG-positive” for Wegener’s Granulomatosis in the original 2010 publication), GPA-negative (“WG-negative” in the original publication), and controls. **(A)** MultiPLIER LV599 is differentially expressed between the three groups (FDR = 1.17e-08). Points and bars represent mean ± 2 * SEM. **(B)** The loadings of the top (highest weight) 50 genes for MultiPLIER LV599 includes ANCA antigens, shown with black bars. The original publication reported differential expression of ANCA antigens in the PBMC fraction.

**Figure S8.**
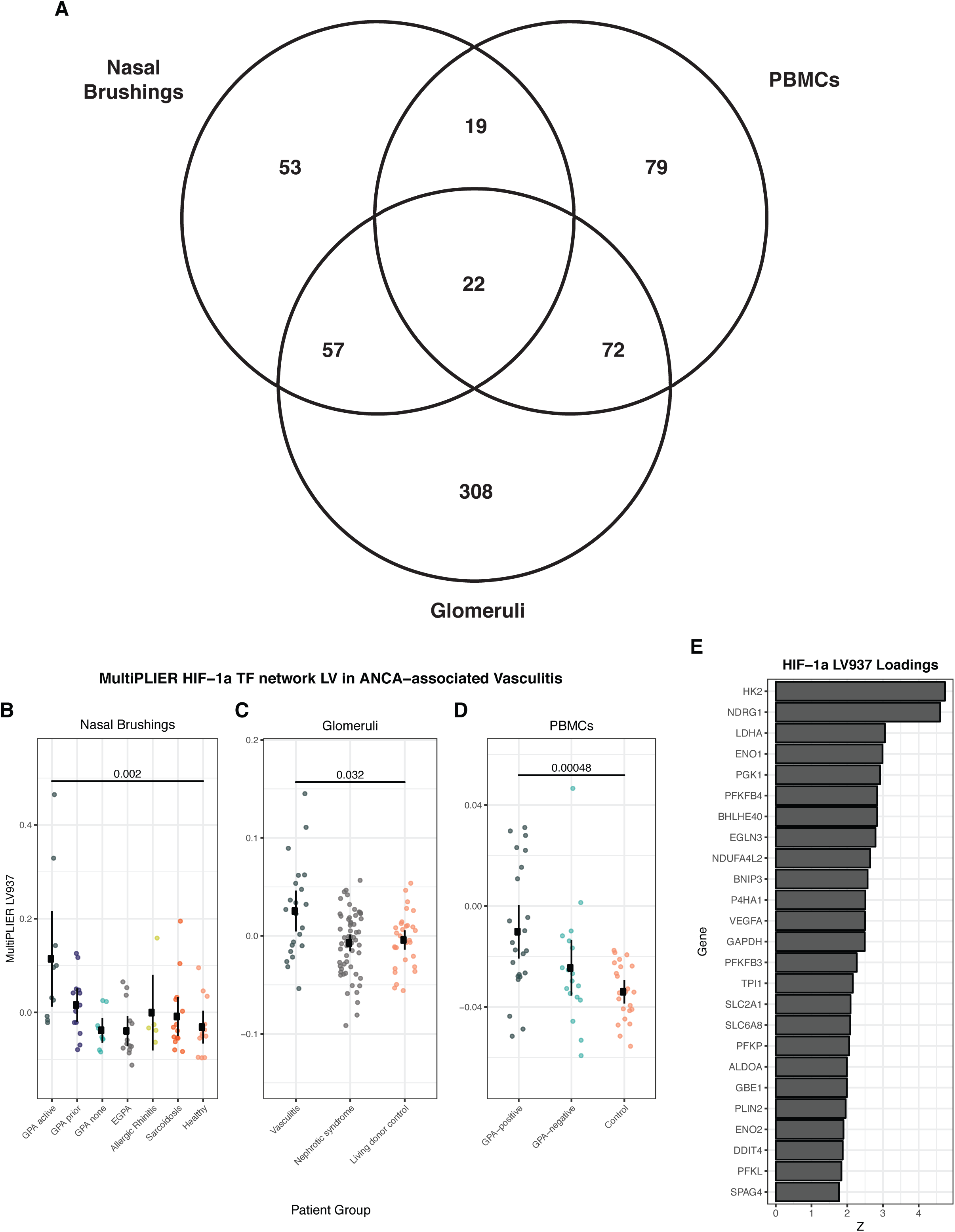
A MultiPLIER-learned latent variable associated with the HIF-1a transcription factor network is differentially expressed in three tissues from ANCA-associated vasculitis (AAV) and shows increased expression in severe or active disease. Related to Figure 7. Differentially expressed latent variables (LVs) were identified by comparing all patient groups and using Benjamini-Hochberg correction (FDR). Latent variables with FDR < 0.05 were considered to be differentially expressed. MultiPLIER LV937 had an FDR < 0.05 in all three cohorts. **(A)** Venn Diagram of latent variables differentially expressed in three tissues. **(B-D)** Jitter plots of MultiPLIER LV937 in three different tissues: nasal brushings (NARES dataset), kidney microdissected glomeruli, and peripheral blood mononuclear cells. Points and bars represent mean ± 2 * SEM. P-values are from a Wilcoxon rank sum test comparing the control group to the AAV group considered to have the most severe or active disease in the cohort. **(E)** The loadings of the top (highest weight) 25 genes for MultiPLIER LV937.

